# Benchmarking cassette-based deep mutagenesis by Golden Gate assembly

**DOI:** 10.1101/2023.04.13.536781

**Authors:** Nicolas Daffern, Irene Francino-Urdaniz, Zachary T. Baumer, Timothy A. Whitehead

**Author notes:** Correspondence to: Timothy A. Whitehead, JSC Biotechnology Building, 3415 Colorado Avenue, Boulder, CO 80305. Authors contributed equally to this work (Nicolas Daffern, Irene Francino-Urdaniz, Zachary T. Baumer).

## Abstract

Protocols for the construction of large, deeply mutagenized protein encoding libraries via Golden Gate assembly of synthetic DNA cassettes employ disparate, system specific methodology. Here we benchmark a broadly applicable Golden Gate method for building user-defined libraries. We demonstrate that a 25 μl reaction, using 40 fmol of input DNA, can generate a library on the order of 1×10^6^ members and that reaction volume or input DNA concentration can be scaled up with no losses in transformation efficiency. Such libraries can be constructed from dsDNA cassettes generated either by degenerate oligonucleotides or oligo pools. We demonstrate its real-world effectiveness by building custom, user-defined libraries on the order of 10^4^ to 10^7^ unique protein encoding variants for two orthogonal protein engineering systems. We include a detailed protocol and provide several general-use destination vectors.

## Introduction

Cassette assembly has become a powerful way to create protein libraries thanks to modern synthesis technologies that can quickly and affordably produce custom DNA fragments^1–4^. Some of the most useful tools for assembling such fragments are Type IIs enzymes, which cut outside of their DNA recognition site and leave a user-defined four base pair overhang. While labs have employed these enzymes to manipulate DNA for the last several decades^5,6^, a major development in their use came in 2008 when Engler *et al*. established a cloning method using the Type IIs enzyme BsaI, which they called Golden Gate assembly^7^. Since then, the method’s popularity has grown^8–10^, due in part to its ability to connect fragments in a specific order and without a restriction scar^11^.

Although the assembly of synthetic DNA by Golden Gate has clear utility in building large protein libraries, a generalized procedure has yet to be established. While several labs have used Golden Gate to build libraries from cDNA and synthetic DNA fragments, their assembly protocols and resulting efficiencies varied widely and often contained time-consuming, system-specific steps ^12–16^. In 2019 Püllmann *et al*. developed a more general Golden Gate protocol for creating site-saturation libraries from oligonucleotides, which was well characterized and included a useful script for primer design^17^. However, the base version of this protocol was only shown to generate a single point mutation and a more complicated protocol involving subcloning was needed to make a library of 60 variants.

Here we provide a simple, broadly applicable Golden Gate procedure that can be used to build large (>10^7^), site-specific, deeply mutagenized libraries. We present data benchmarking the procedure’s efficiency under different use conditions and demonstrate its effectiveness in constructing libraries from mixed base-containing oligonucleotides and custom synthesized oligo pools. We also provide a detailed protocol with discussion of important design considerations and three general-use destination vectors deposited on Addgene.

## Results/Discussion

### A standardized protocol for library generation

The general workflow for library generation using our protocol involves design and creation of a destination vector and mutagenic cassettes, assembly via a Golden Gate reaction, and transformation into bacteria. The destination vector must be designed and created with a selection marker, BsaI sites, and specific overhangs such that cassette(s) introduction results in reconstitution of a full-length gene encoding sequence, and replacement of the marker which allows rapid assessment of incorporation efficiencies (**Figure 1A**). In parallel, cassettes are designed with BsaI sites and overhangs arranged for sequential insertion into the destination vector (**Figure 1A**) using subsets of the high-fidelity overhangs outlined by Potapov *et al*. to reduce inefficiencies from imperfect ligation^18^. Subsequently, the destination vector and cassettes are mixed with BsaI-HFv2 and T4 ligase and PCR cycled between 37 °C and 16 °C, during which time the selection marker and cassette ends are removed, the cassettes anneal with the vector, and the annealed DNA is ligated.

**Figure 1.**
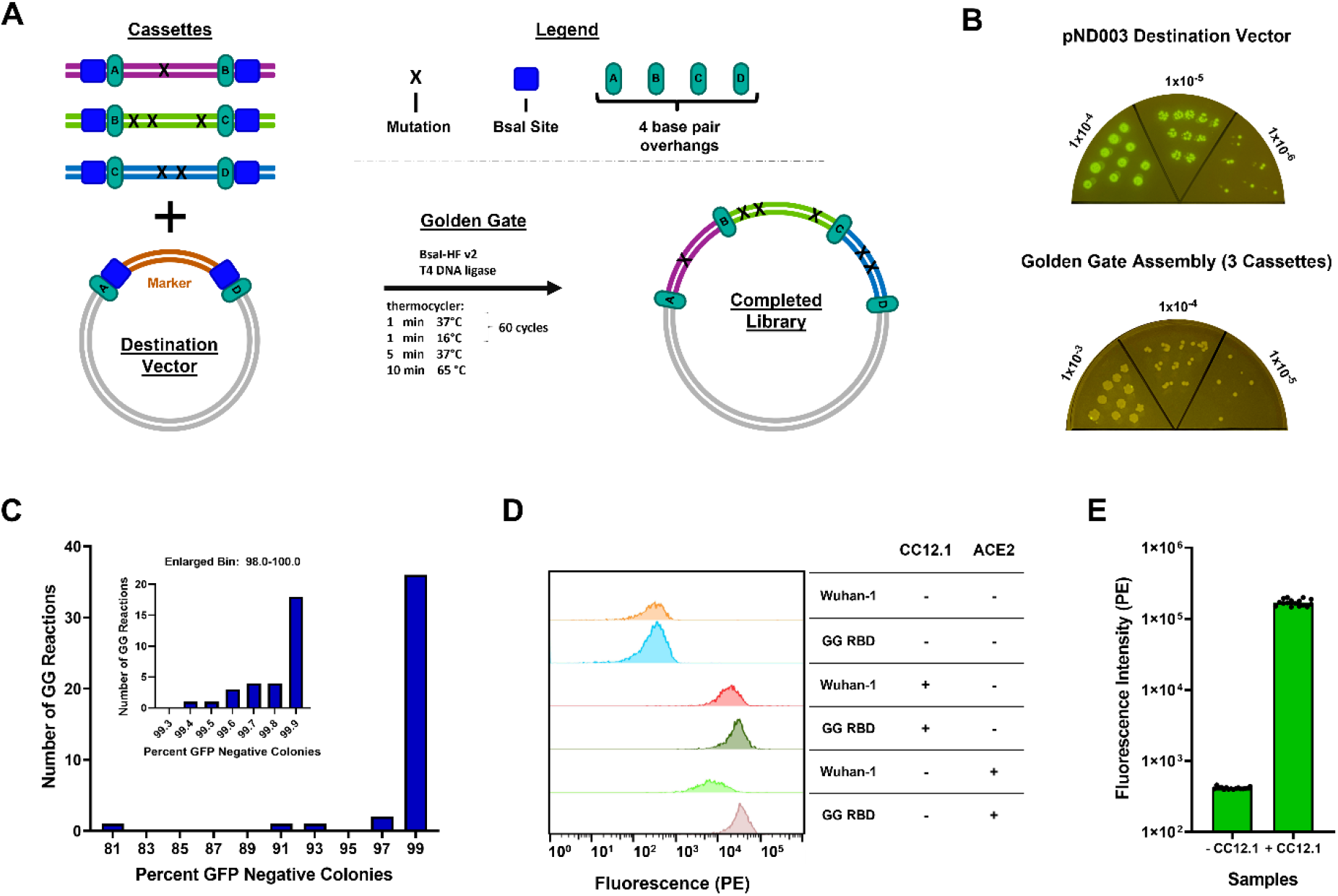
Establishing the functionality of a Golden Gate protocol for library generation. **A**. Schematic showing library generation by assembly of mutagenic cassettes into a destination vector via Golden Gate. **B**. Dilution plating of E.coli transformations with 40 fmol of our destination vector pND003 (left) and a Golden Gate reaction performed with 40 fmol of both pND003 and three wild type PYR1 cassettes (right). Listed numbers represent fold dilution. **C**. Histogram showing the percent cassette incorporation for all performed Golden Gate reactions. Inset shows all reactions from the 98-100% bin redistributed in bins with 0.1% intervals. Incorporation percentages were calculated by comparing the number of green (GFP) and white (non-GFP) colonies on dilution plates after transformation. **D**. Functional comparison of GG RBD and Wuhan-1 RBD using yeast surface display. RBD displaying yeast were incubated in the absence (-) or presence (+) of saturating concentrations of PE-labeled CC12.1 (antibody) or ACE2, followed by assessment of binding by flow cytometry. **E**. Assessment of CC12.1 binding for 16 colonies displaying RBD from 4 different Golden Gate reactions. Individual values represent mean PE fluorescence intensity of RBD-displaying populations.

We developed a modified Golden Gate protocol that allowed us to rapidly assemble libraries while maintaining high numbers of transformants. For the base version of this protocol, we used a 25 μl reaction containing 40 fmol of the destination vector and each cassette. Additionally, our reactions were PCR cycled for 60 cycles with each step only lasting one minute, as described by Strawn *et al*.^19^, allowing the Golden Gate reaction, and bacterial transformation to be performed in under 8 hours (**Figure 1A**). We have included a protocol describing the design and creation of libraries using this technique in more detail (**Supporting Information**). Using this protocol, with properly designed cassettes and a destination vector in hand, a new library for any protein can be generated in a single day. To facilitate this, we built general use destination vectors for yeast surface display (pND003), MBP-tagged protein expression in *E. coli* (pND004), and yeast two-hybrid assays (pND005), each of which contain a GFP marker and BsaI sites with high fidelity overhangs. To confirm their functionality, a Golden Gate reaction was performed with pND003 and cassettes that encode for a monomeric variant of the plant abscisic acid receptor PYR1 (H60P, N90S)^20^. Comparison of plates from a transformation of pND003 alone and a transformation of the Golden Gate reaction confirmed cassette insertion and demonstrated that removal of the GFP marker allowed for assessment of incorporation efficiencies (**Figure 1B**).

We used numbers of GFP negative colonies to calculate incorporation percentages for all Golden Gate reactions described in this paper with a median incorporation of 99.7% (range 81% to >99.9%, n=40) (**Figure 1C; Supplemental Table 1)**.

To verify that our protocol would result in genes encoding full-length, functional protein, we assessed the ability of an SARS-CoV-2 Omicron chimeric S RBD to bind ACE2 and the neutralizing monoclonal antibody CC12.1^21,22^. For these experiments we created a destination vector with the coding sequence for the Omicron chimeric RBD (SARS-CoV-2 S RBD (333-541) Wuhan-1 with mutations S477N, E484A, Q498R, N501Y and Y505H) and the corresponding dsDNA coding cassettes, assembled them into the RBD destination vector using our Golden Gate protocol, transformed the resulting plasmids into yeast, and expressed the isogenic Golden Gate-derived RBDs (GG RBD) on the surface of yeast. As a positive control, we also expressed the previously described Wuhan-1 S RBD N343Q (Wuhan-1)^22^ on the surface of yeast. We then compared the ability of the GG RBD and Wuhan-1 to bind Fc-ACE2 and CC12.1 (which contains an Fc) using flow cytometry. When labeled with an anti-Fc PE, we observed specific PE fluorescence resulting from binding to ACE2 or CC12.1 for both GG RBD and Wuhan-1 (**Figure 1D**). We then tested GG RBD functionality in a similar manner for 16 different yeast colonies from four separate Golden Gate reactions and saw consistent levels of CC12.1 binding for all 16 variants, highlighting the ability of this protocol to reproducibly assemble full-length sequences (**Figure 1E**).

### Benchmarking a scalable single day Golden Gate library generation protocol

Next, we benchmarked the number of transformants that can be generated using this protocol as a function of cassette number, reaction size, and input DNA. For all these experiments we used our pND003 vector and PYR1 cassettes. Our baseline for all three experiments was a 25 μl reaction containing 40 fmol of the destination vector and a single PYR1 cassette, which generally results in 7-8×10^5^ transformants and >96% cassette incorporation (**Figure 2A-C, Supplemental Table 1**).

**Figure 2.**
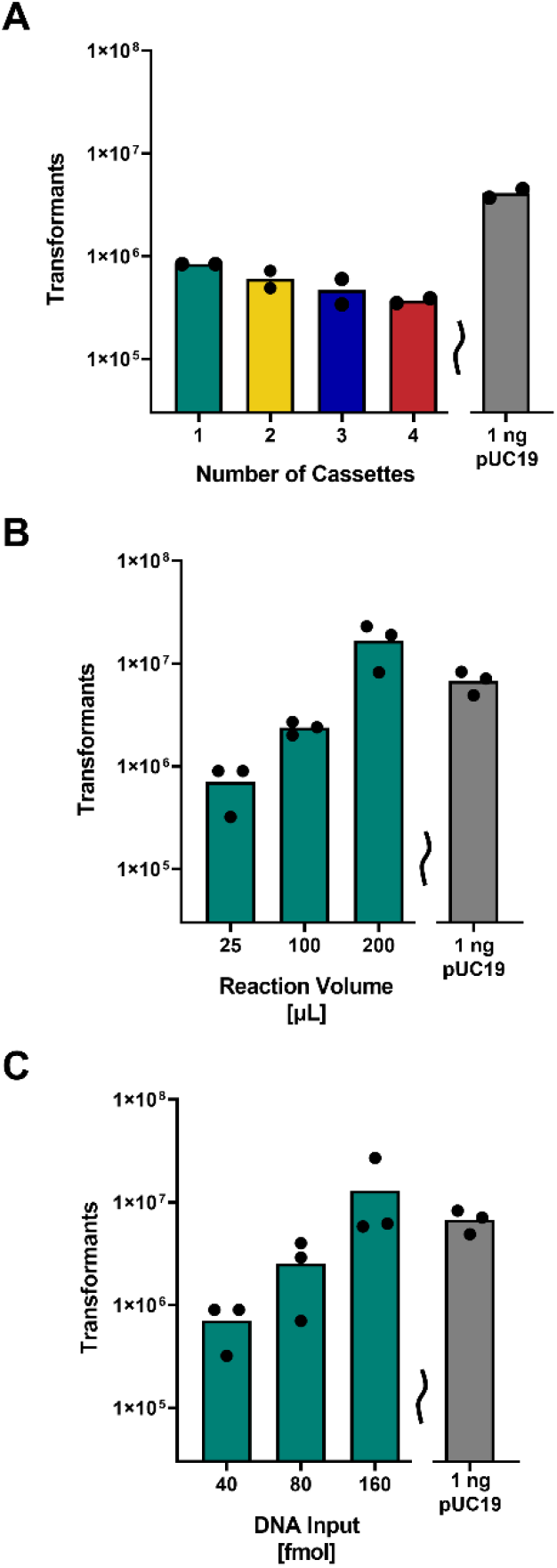
Benchmarking a Golden Gate protocol for library generation. **A**. Transformation efficiencies for Golden Gate reactions performed with differing numbers of cassettes. 25 μl reactions were performed in duplicate with 40 fmol of both pND003 and one, two, three, or four wild type PYR1 cassettes. Numbers of transformants were calculated by plating serial dilutions of the recovered cells. pUC19 is shown as a control reaction for assessing efficiency of electrocompetent cells. **B**.,**C**.: Transformation efficiencies as a function of Golden Gate reaction size (**B)** or input DNA (**C**). A baseline 25 μl reaction using 40 fmol of both pND003 and one wild type PYR1 cassette was compared with reactions having increased size or input DNA. Reactions were performed in triplicate. Numbers of transformants were calculated by plating serial dilutions of the recovered cells. (**B**.) All reaction components were scaled 1:1, including DNA concentration, with increasing volume. (**C.)** The total amount of input DNA for the destination vector and cassette were increased while all other reaction conditions were held constant.

Since previous studies have noted decreasing Golden Gate efficiencies when increasing the number of ‘parts’ (number of cassettes plus destination vector)^13,23^, we sought to quantitate the number of transformants as a function of the number of input cassettes. For this experiment, we performed four reactions with one to four PYR1 cassettes under conditions that were otherwise identical to the baseline reaction. We found that transformation efficiencies were modestly affected by increasing cassette number, with a four-cassette (five-part) assembly showing a minor reduction in transformants (3.7×10^5^, n = 2) over a one-cassette (two-part) assembly (8.4×10^5^, n = 2) (**Figure 2A, Supplemental Table 1**). Thus, increasing cassette numbers should not hinder most library designs.

We then assessed the ability of our protocol to be scaled by total reaction size by comparing our baseline 25 μl reaction with 100 μl and 200 μl reactions. We found that the number of transformants scaled at least linearly, with the 200 μl reaction (8x volume) generating an approx. 24-fold increase in transformants with no loss in incorporation efficiency (p-value 0.03**; Figure 2B, Supplemental Table 1**). We speculate that this trend results from decreased relative DNA loss while working with small constant volumes during the PCR cleanup and transformation steps. Since a 100 μl reaction can be performed in a single PCR tube, several orders of magnitude higher numbers of transformants should easily be achieved by pooling multiple Golden Gate reactions in a single PCR cleanup column.

Subsequently we tested scaling of DNA concentration by comparing our baseline reaction with reactions in which the amount of destination vector and cassette DNA was increased to 80 and 160 fmol, without increasing reaction size. We again observed a linear increase in transformants, with 160 fmol (4×increase) of input DNA resulting in a 19-fold increase in transformants (p value 0.13 for whether the 160 fmol reaction gives more than a 4× increase in number of transformants) and no loss in incorporation efficiency (**Figure 2C, Supplemental Table 1**). Thus, reactions can be scaled by both volume and DNA concentration.

### Creation of a site-specific protein libraries using both DNA Ultramers and oligo pools

We next assessed the ability of our protocol to produce complete libraries from different types of synthetic DNA by building libraries for the Omicron chimeric RBD. For this we designed three site-specific combinatorial libraries, all covering the same 110 contiguous positions, assembled using a combination of three mutant cassettes (**Figure 3A**).

**Figure 3.**
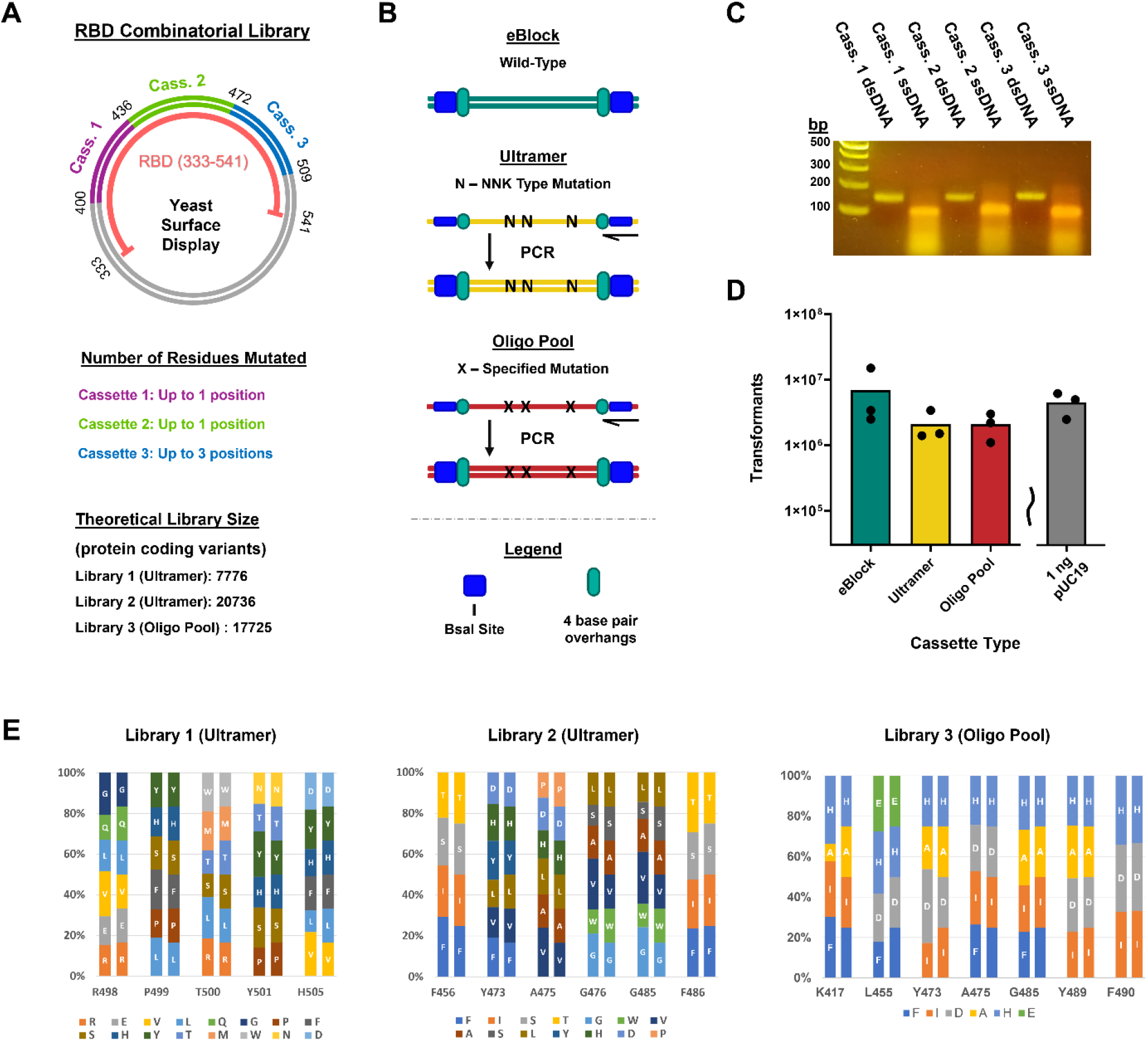
Golden Gate assembly of SARS-CoV-2 S RBD deep mutational libraries. **A**. Schematic of the assembled plasmid for the S RBD combinatorial library. Unmutated S RBD residues 333-399 and 510-541 are encoded in the destination vector pIFU037 while residues 400-509 are encoded by three mutagenic cassettes. Cassettes can code for wild-type or mutant residues at each mutational site **B**. Different cassette DNA inputs for the Golden Gate assembly. eBlocks (dsDNA) are used as is, while Ultramers (mixed base-containing long oligonucleotides) and oligo pools are obtained as lyophilized ssDNA and require PCR synthesis using a reverse primer to generate dsDNA. **C**. Gel electrophoresis of three RBD library Ultramer cassettes before (ssDNA) and after PCR (dsDNA). Each cassette is 170 nts. **D**. Transformation efficiencies for Golden Gate reactions performed with pIFU037 and the three RBD library cassettes, each generated from different types of DNA. 25 μl reactions were performed in triplicate with 40 fmol of destination vector and cassettes. Numbers of transformants were calculated by plating serial dilutions of the recovered cells. pUC19 is shown as a control reaction for assessing efficiency of electrocompetent cells. **E**. Mutational distributions for RBD libraries. Expected (right bars) vs observed mutational frequency (left bars) at each mutated residue based on deep sequencing.

We generated cassettes from synthetic dsDNA (eBlocks), mixed-base degenerate long oligonucleotides (Ultramers), and ssDNA sourced from custom oligo pools (**Figure 3B**). Generally, we used eBlocks to encode unmutated regions of protein, while Ultramers and oligo pools were used as mutagenic cassettes. dsDNA cassettes are generated from single-stranded Ultramers or oligo pool DNA using PCR with a reverse primer (**Figure 3B**). For these experiments, we assembled Libraries 1 and 2 from PCR-amplified mutant Ultramer cassettes and Library 3 from PCR-amplified mutant oligo pool cassettes (an example of this amplification for Library 1 is shown in **Figure 3C**). For comparison, we performed a fourth assembly with eBlock cassettes containing no sequence variation.

We performed PCR amplification of the oligo pools and Ultramers as well as the Golden Gate reactions using the base method described in our protocol, except for increasing input DNA from 40 to 200 fmol. We found similar transformation efficiencies using Ultramers and oligo pools, with both resulting in approximately 1.5×10^6^ transformants, and a slightly higher efficiency with eBlocks, which resulted in approximately 7.0×10^6^ transformants (**Figure 3D**). Thus, our protocol generates consistently high numbers of transformants with different types of input DNA, giving the user flexibility when designing their libraries.

To assess library quality, completed libraries were deep sequenced at a depth ranging from 5.9e5 to 2.9e6 reads (**Supplemental Table 2**). In contrast to other user-defined mutagenic protocols with high wild-type sequence carryover^24^, no library contained more than 0.2% wild-type reads, and all libraries contained at least 96.7% of the desired variants (96.7, 99.9, 99.9%). The libraries ranged from 80.9-86.8% of on-target sequences. 9% of the oligo pool-derived library coded for chimera sequences that contained unintended combinations of designed mutations. These could have arisen during our deep sequencing preparation, as chimera formation is known to occur at low abundances during the associated PCR steps^22^. Alternatively, chimera formation could occur during the cassette generation step of our protocol; our analysis is unable to distinguish between these possibilities. We assessed library uniformity by comparing the theoretical frequency of different residues at each mutated position with the observed frequency seen in our deep sequencing data (**Figure 3E**). The relative frequencies observed varied between 0.34-1.52-fold as compared with expectations (n=62). Together, this data demonstrates that our Golden Gate protocol can generate user-defined combinatorial mutational libraries with almost complete coverage, little to no wild-type carryover, and near-uniform individual mutational distributions.

### Creation of a large site-specific mutational library

We next assessed if our protocol could be scaled to generate larger, high-quality libraries by designing a library for T7 RNA polymerase coding for 1.1×10^7^ theoretical protein variants using a reduced codon alphabet^25^ (**Figure 4A**). We chose to build the library using Ultramer-based cassettes, where mutations spanned 363 nucleotide bases (V725-F845) and were grouped at three locations across two cassettes (**Figure 4A**). The second cassette, containing mutations at positions 781-786 and 845, was designed using two overlapping Ultramers (**Figure 4A, 4B**). To construct the library, we performed two separate 200 μl Golden Gate reactions that each contained 200 fmol of destination vector and each cassette, and pooled the DNA for use in a single transformation, generating 5.6×10^7^ transformants with an incorporation efficiency of >99% (**Figure 4C**).

**Figure 4.**
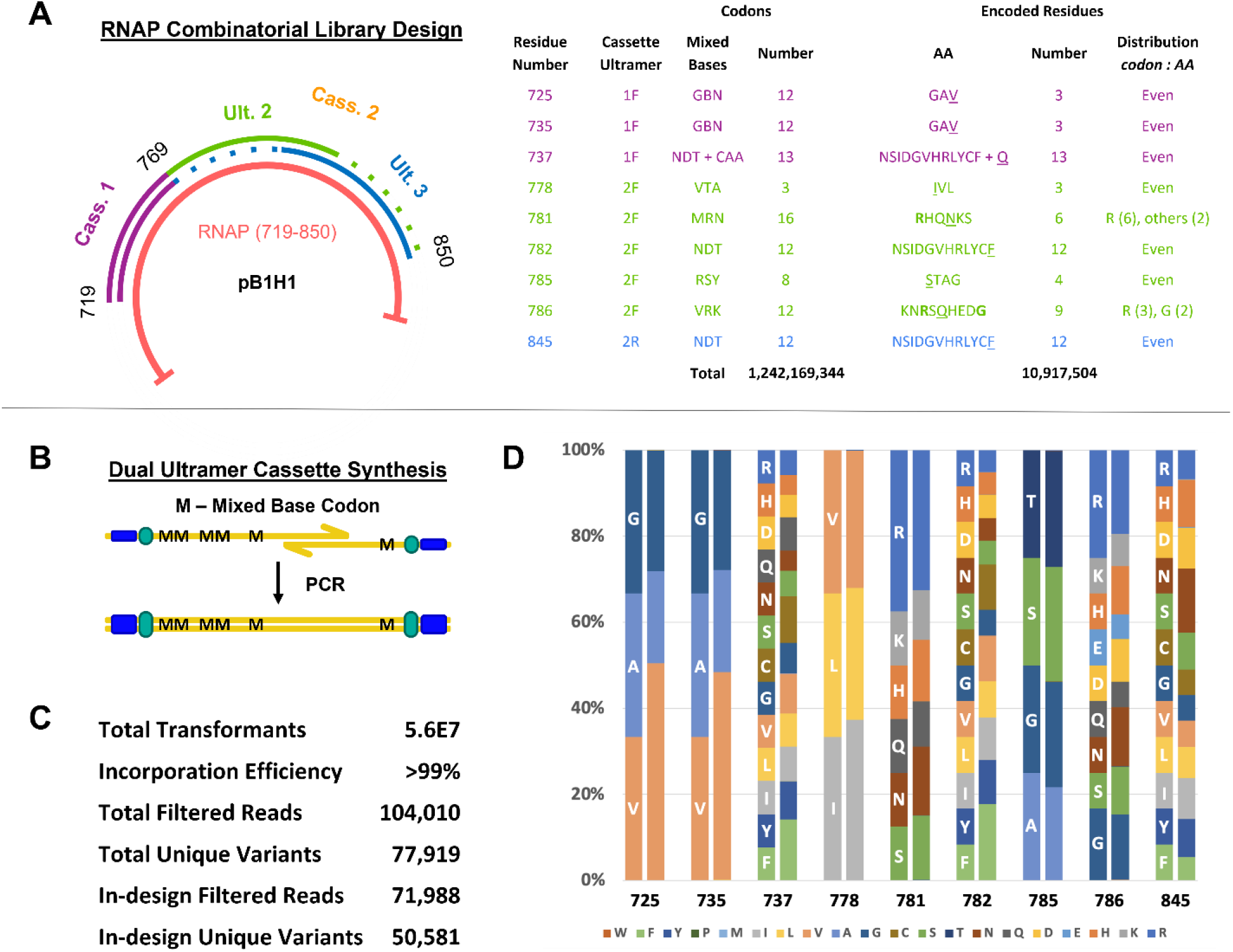
Golden Gate assembly of a large T7 RNAP library. **A**. Schematic of the assembly strategy and mutational designs for the T7 RNAP library. The assembly used two cassettes: one standard cassette generated from a single Ultramer, and one long cassette generated from two overlapping Ultramers (left). Different sets of mixed base codons were used to code for subsets of amino acids at each mutational site (right). Underlined single letter amino acids are the wild type residue at a given position and bold residues are encoded by a greater relative number of codons as a function of the mixed bases chosen. **B**. Schematic for using two overlapping Ultramers as forward and reverse primers to generate a longer cassette. **C**. Summary Library Statistics. Filter conditions included correct length of merged reads, the presence of “N” in the merged reads and the presence of stop codons. In-design reads exclude any mutations at protein level which are not in the design space. **D**. Expected (left bars, with single letter amino acid code) vs observed mutational frequency (right bars) at each library residue. The sum of all amino acids not in the set of designed mutations was less than 0.5% at each position.

Subsequently, we performed limited deep sequencing to assess the quality of our library. We obtained 104,010 reads after quality filtering and observed 77,919 total unique protein encoding variants, including 50,581 in-design protein encoding variants, with no wild-type sequences, further demonstrating the ability of our protocol to limit wild-type carryover. To assess library uniformity, we compared the theoretical frequency of different amino acids at each mutational position with the observed frequency seen in our deep sequencing data (**Figure 4D**). The relative frequencies observed varied between 0.6-2.1-fold compared with expectations (n=65), showing that large libraries can be assembled with minimal mutational bias.

## Conclusions

Although the need for large, site-specific libraries for protein engineering workflows is almost universal, the methods for generating such libraries vary greatly. Modern DNA synthesis technology, which allows for the rapid creation of short DNA fragments containing user-defined mutations, has the potential to streamline these methods and make libraries more consistent and affordable. Here we provide a simple, broadly applicable protocol detailing how to build large libraries from synthetic DNA cassettes using Golden Gate assembly. To provide an accurate assessment of this protocol’s capabilities, we benchmarked the obtainable transformation and incorporation efficiencies as a function of cassette number, reaction size, and input DNA. We then demonstrated its real-world applicability by creating libraries of different sizes in two orthogonal protein engineering systems. First, we generated libraries for the SARS-CoV-2 S RBD on the order of 10^4^ members and demonstrated by deep sequencing that near-uniform, near-complete libraries can be made equally well using mixed base oligonucleotides or oligo pools. Subsequently, we built and characterized a library encoding 1.1× 10^7^ theoretical T7 RNAP to show that this technique is broadly applicable and that uniformity and coverage are maintained when our protocol is scaled up. We expect our benchmarking will facilitate implementation of cassette-based Golden Gate mutagenesis for the protein engineering and design community.

## Methods

### Construction of destination vectors and cassettes

Destination vectors were created by combining a source vector and a GFP-encoding insert from pYTK047^10^ or pEDA5^24^ using Gibson Assembly^26^ or restriction enzyme cloning. Oligo pool and Ultramer derived double-stranded cassettes were obtained by performing PCR with single-stranded source DNA and a single reverse primer, followed by gel extraction. Sequences for primers, completed destination vectors, and wild-type versions of cassettes are listed in the Supporting Data (**Supplemental Table 2, Supplemental Data**).

### Performance and assessment of Golden Gate reactions

Details for the performance and assessment of Golden Gate reactions are listed in the Supporting Data (**Supplemental Methods**). In brief, Golden Gate reactions were performed with a standard protocol that used 40 fmol of each input DNA and was temperature cycled 60 times with 1-minute steps to reduce waiting times. Alterations to this protocol are noted in the text. Reactions were then column purified and transformed into electrocompetent cells, along with a pUC19 control. Transformation and incorporation efficiencies were determined by counting the number of white and green cells after serial dilution plating.

### Characterization of Golden Gate assembled RBD constructs

Binding of GG and Wuhan-1 RBDs to ACE2/CC12.1 was assessed by yeast display as described by Francino-Urdaniz *et al*^22^.

### DNA deep sequencing

Libraries were prepared for deep sequencing following the “Method B” protocol from Kowalsky *et al*^27^. PCR protocols and primers used in library preparation and a detailed description of the deep sequencing data analysis are provided in the Supporting Data (**Supplemental Methods, Table S3**).

## Author Information

### Author Contributions

Conceptualization: T.A.W., N.D., I.F.U., and Z.T.B. Experimental Design: T.A.W., N.D., I.F.U., and Z.T.B. Performed Experiments: N.D., I.F.U., and Z.T.B. Data Analysis: T.A.W., N.D., I.F.U., and Z.T.B. Wrote First Draft: N.D. Writing & Editing: T.A.W., N.D., I.F.U., and Z.T.B. Supervision: T.A.W. Funding Acquisition: T.A.W. and Z.T.B.

**Notes**

The authors declare no competing financial interest.

## Acknowledgements

This work was supported by the National Institute Of Allergy And Infectious Diseases of the National Institutes of Health (Award Numbers 1R21AI174157-01 and 5R01AI141452-05 to T.A.W.), the Defense Advanced Research Projects Agency Advanced Plant Technologies (grant HR001118C0137 to T.A.W. - Approved for Public Release, Distribution Unlimited), the National Science Foundation (grant NSF CBET-2030221 to T.A.W.), and the National Science Foundation Graduate Research Fellowship Program (Z.T.B. DGE Award Number 2040434, fellow ID: 2021324468).

## Data Availability

The plasmids pNDD003, pND004, and pND005 and their corresponding sequences have been deposited at AddGene (to be released upon publication). All raw deep sequencing reads have been deposited at the Sequencing Read Archive (SRA to be updated upon publication). Raw data used to prepare figures 1C, 2A-C, and 3D are present in **Supplemental Table 1**.

## Supporting Information

The following Supporting Information is available free of charge at the <u>ACS website</u>:

Supplemental methods; Table of conditions used in all Golden Gate reactions; Table of all primer sequences used; Detailed protocol for performing Golden Gate cassette mutagenesis; Complete sequences of all plasmids and cassettes used

