## Supporting Information for "Benchmarking cassette-based deep mutagenesis by Golden Gate assembly"

[‡] Authors contributed equally to this work

***This supplementary information contains:***

***Supplemental Note 1:*** Detailed Golden Gate-based cassette mutagenesis protocol

***Supplemental Methods***

***Supplemental Table 1:*** Table of all Golden Gate reactions performed

***Supplemental Table 2:*** Library statistics for S RBD sequenced libraries

***Supplemental Table 3:*** Primer sequences

***Supplemental Data:*** Plasmid and cassette sequences.

### *Supplemental Note 1: User-defined library generation by Golden Gate assembly*

#### BACKGROUND & PROTOCOL OVERVIEW

Golden Gate assembly is a one-pot DNA assembly technique that uses temperature cycling with a type IIS restriction enzyme (BsaI-HFv2 reported here), T4 ligase, and input DNA to clone multiple DNA fragments sequentially into a vector. Generally, for this protocol, a PCR thermocycler cycles between a 37 °C step and a 16 °C step. During the 37 °C step BsaI generates four base pair overhangs downstream of its GGTCTC(X) binding motif and during the 16 °C step the reannealed DNA is ligated by T4 ligase. BsaI sites on the DNA fragments are designed such that overhangs will match complementary overhangs on upstream and downstream fragments or vector ends, allowing the fragments to be assembled in order. Additionally, the orientation of the BsaI sites on the vector and fragments are such that they are removed from the final product. Thus, cycling of these cutting and ligating steps continues to act on the input DNA and leaves the final product untouched, resulting in high incorporation efficiencies.

Here we detail a version of this protocol that can be applied to any vector and protein coding sequence to generate large, site-specific, combinatorial libraries by assembling synthetic mutagenic DNA fragments that we denote cassettes into a destination vector. Cassettes are linear dsDNA and can be purchased as synthetic dsDNA (e.g. gBlocks/eBlocks from IDT) or assembled from oligonucleotide or oligo pool ssDNA. Destination vectors have some, or all, of the protein coding sequence replaced with a selective marker, such that mutated cassettes coding for that sequence replace the marker during the Golden Gate reaction, resulting in functional plasmids. Users can follow our protocol to make their own destination vector or use one of our premade vectors; our protocol provides a detailed explanation for designing cassettes that work with pND003, pND004, and pND005. After the destination vector is obtained, cassette(s) (one to four demonstrated) are generated from ssDNA by PCR and co-incubated with the destination vector in a Golden Gate reaction to assemble the library, which is then transformed into *E. coli*. We describe a streamlined protocol such that, once the destination vector is in hand, new libraries can be built in a single day.

The protocol is listed in three sections: (1.) Design and construction of the destination vector; (2.) Design and construction of double stranded cassette DNA; (3.) Golden Gate assembly and transformation into *E. coli*. Additionally, we provide examples for different steps based on the creation of the SARS-CoV-2 S RBD (333-541) yeast display libraries (RBD libraries).

#### MATERIALS AND SUPPLIES

*All customized DNA fragments were purchased from IDT

| **Item** | **Supplier** | **Catalog number** |
|---|---|---|

| | | |
|---|---|---|
| **Section 1: Construction of destination vector** | | |
| pETconNK | Klesmith et al. 2017[1] | https://www.addgene.org/81169/ |
| knock-out gene | Lee et al. 2015[2] | https://www.addgene.org/65154/ |
| **Section 2: Construction of dsDNA cassettes** | | |
| Cassette 1 ultramer/opool | This study | Table S4 |
| Cassette 2 ultramer/opool | This study | Table S4 |
| Cassette 3 ultramer/opool | This study | Table S4 |
| Forward primer | This study | Table S5 |
| Q5 High-Fidelity 2X Master Mix | NEB | M0492S |
| KAPA HiFi Ready Mix | Roche/Fisher | 7958927001 |
| Nuclease Free Water | IDT | 11-05-01-14 |
| Ultrapure Agarose | Invitrogen | 16500-500 |
| TAE Buffer (Tris-acetate-EDTA) (50X) | Thermo Scientific | B49 |
| SYBR Safe DNA gel stain | Invitrogen | S33102 |
| Gel loading dye, Purple | NEB | B7024 |
| Nucleo-Spin® Gel and PCR Clean-up kit | Macherey-Nagel | 740609.250 |
| **Section 3: Golden Gate assembly and transformation into *E. coli*** | | |
| Destination vector | This study | Step 1 |
| dsDNA Cassette 1 | This study | Step 2 |
| dsDNA Cassette 2 | This study | Step 2 |

| | | |
|---|---|---|
| dsDNA Cassette 3 | This study | Step 2 |
| BsaI-HF v2 | NEB | R3733S |
| T4 DNA ligase | NEB | M0202S |
| T4 DNA ligase reaction buffer | NEB | B0202S |
| Nuclease Free Water | IDT | 11-05-01-14 |
| XL1-Blue Competent Cells | Agilent | 200249 |
| TransforMax EC100 Competent Cells | Biosearch Technologies | EC10010 |
| Corning® Square Bioassay dish | Sigma | CLS431272 |
| Glass Beads | Sigma | G8772 |
| Kanamycin | GoldBio | K-120-25 |
| ZymoPURE II Plasmid Midiprep kit | Zymo Research | D4201 |

**PROTOCOL**

Section 1: Design and construction of the destination vector

**Background:** We built three destination vectors (MBP *E. coli* expression, yeast surface display, yeast two-hybrid), each containing a GFP marker and BsaI recognition sequences next to high-fidelity overhangs **(Supplemental Note, Figure 1)**. If any of these destination vectors are suitable, section 1 can be skipped.

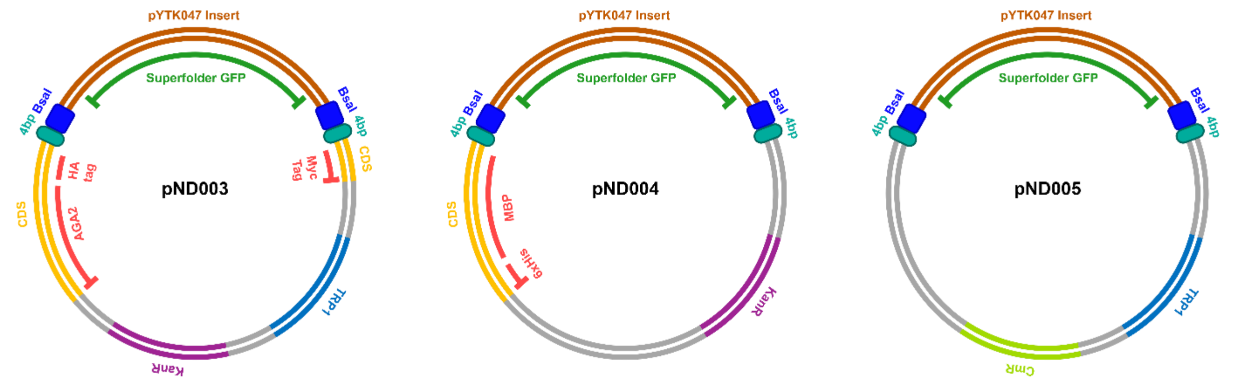


***Supplemental Note, Figure 1.*** *Premade general-use destination vectors for Golden Gate-based cassette mutagenesis*

If these destination vectors are unsuitable, you can create your own destination vector. The following describes how to make a destination vector using Gibson Assembly, although a variety of cloning techniques could be used.

**1: Pick a backbone**

Pick a backbone for your destination vector which contains the protein of interest and the desired vector components (tags, promoters, selection markers, etc). Check to see if the vector already contains BsaI sites. If so, these should be removed by site-directed mutagenesis before proceeding.

**2: Choose a selection marker**

Our lab has used both GFP and RFP but other selection markers, such as LacZ[3,4], LacI[4], β-galactosidase[4] or sacB[5], have been demonstrated to be effective for GG assembly.

**3: Choose the start and stop points for your cassettes**

Before picking the start and end points for your library, you must choose the general region covered by your cassettes. Here you have two options. If you already have your protein in the desired vector, your cassettes can cover only the coding sequence containing your mutation sites. This allows for the fewest number of cassettes, is the simplest to design, and is ideal for large protein-coding sequences but requires that a new destination vector be created if you mutate a different region of your protein. Alternatively, you can have your cassettes begin and end outside the protein coding region, such that any library can be made for any protein without building a new destination vector (as with our premade destination vectors).

Once you have chosen the region covered by your cassettes, pick a set of high-fidelity 4 base pair overhangs from Table 1 in Potapov *et al*[4]. Then find a four base pair sequence near the beginning and end of that region from that set. These will be the start and end points for your cassettes.

#### 4: Design primers for creation of the destination vector

Design primers to clone the desired selective marker and BsaI sites into the destination vector, in place of the backbone sequence covered by the cassettes. If using Gibson Assembly, this includes two primers to amplify the selection marker from its source as well as two primers to amplify the backbone vector. The primer set for amplifying the backbone should add BsaI sites directly adjacent to the first and last overhangs and 20 base pairs of homology with the selection marker from each end of the PCR fragment. Importantly, the BsaI sites should be in between the overhang and selection marker sequence (**Supplemental Note, Figure 2**).

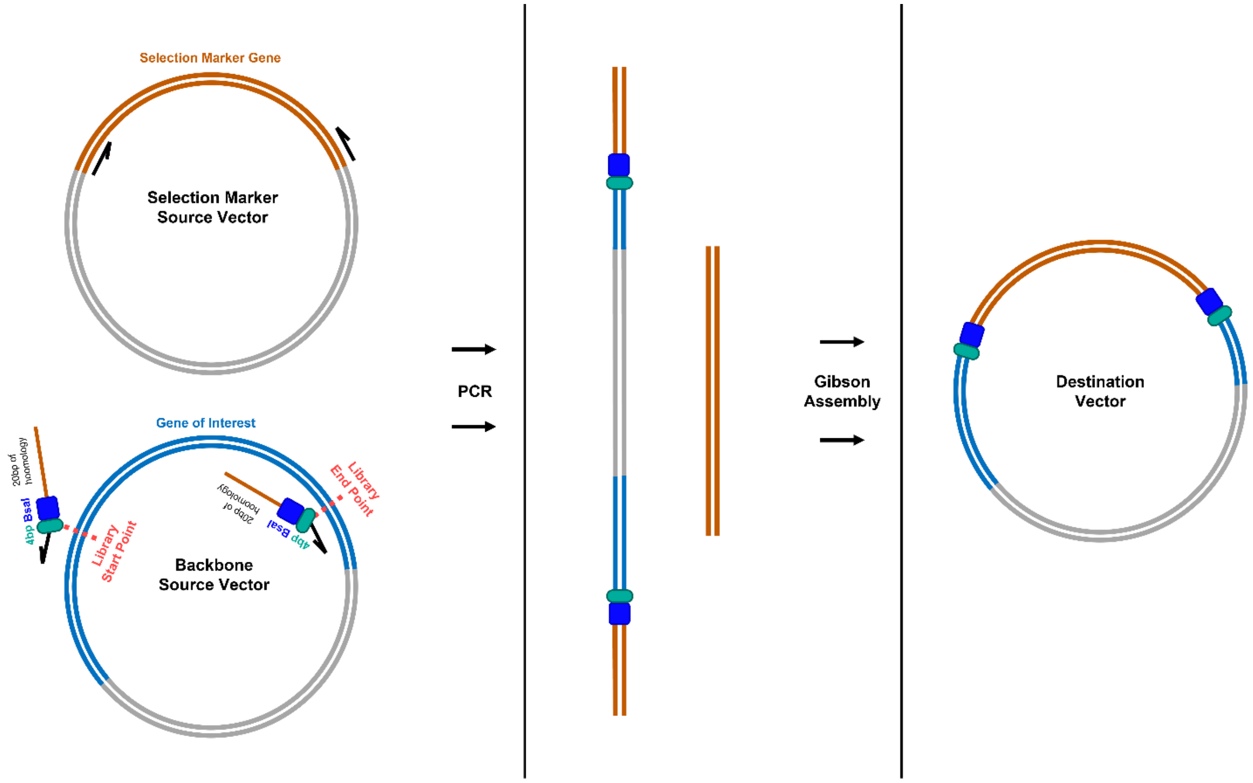


***Supplemental Note, Figure 2.*** *Building a destination vector using Gibson Assembly*

#### 4. Create the destination vector

If using Gibson Assembly, the template vector and selection marker insert should be amplified using a standard PCR reaction. The resulting PCR products are then gel purified and combined by Gibson Assembly[6].

Section 2: Design and construction of cassettes

**Background:** After designing a destination vector or choosing to use one of our premade vectors, you must design cassettes that contain your mutated protein-coding sequence, high-fidelity overhangs, and a BsaI recognition sequence. Additional sequences must be included when designing cassettes for use with pND003, pND004, or pND005 (described below). These cassettes are ordered as synthetic ssDNA fragments and are converted to dsDNA using a reverse primer.

**1: Cassette designs**

The first step in designing cassettes is choosing your library's start and end points with respect to your protein coding sequence. If you created your own destination vector, as per section 1, these have already been chosen. If you are using our premade vectors, these can be chosen arbitrarily. Next, you must decide the number and length of your cassettes, which is usually based on the maximum length of the synthetic DNA fragments you are using (It is important to remember that the length of each fragment must include at least 7 base pairs on either side of the coding sequencing for the BsaI site). This will then determine the general areas in your coding sequence where each cassette will begin and end.

Within these areas, look for four base pair sequences that match high-fidelity overhangs from the set you used for the destination vector or that are listed for our premade vectors at the bottom of **Supplemental Note, Table 1**. Note that all overhangs must be from the same set, cannot be used more than once and that residues coded for by base pairs in the overhangs cannot be mutated. These overhang sequences will be the starting and ending points of each cassette. If designed correctly, the first and last four protein-coding base pairs of each cassette will match the first or last four protein-coding base pairs of the adjacent cassettes (see Example 1). Next, you will add BsaI recognition sequences directly adjacent to the overhangs **(Supplemental Note, Figure 3)**. Note, it is a good idea to check that any mutations you make do not introduce a BsaI site into your coding sequence. We also added short sequences outside of the BsaI site, if no other sequence was present, to ensure BsaI binding; we have not tested whether this is necessary to achieve high transformation efficiencies.

| | | | |
|---|---|---|---|
| Short sequence we included to support binding of BsaI to its recognition sequence. Removing it may not affect efficiencies but has not tested by us. | GCCGT | Only needed when working with our premade destination vectors or designing destination vectors with cassettes that extend beyond the protein coding sequence. Constant sequences that code for promoters, stop codons, terminators, and tags in source vectors. Exact sequences that match our premade vectors are found in **Supplemental Table 3** (colored in red)**.** | NNNNNNNNNN |
| BsaI recognition sequences (forward/ reverse). | GGTCTCA/TGAGACC | User's protein coding sequence. X represents the first and last mutation sites in the cassette | NNXN. . . . . . . .NXNN |
| 4 base pair high-fidelity overhang sequence. When working with our premade destination vectors, the initial overhang in the first fragment and final overhang in the last fragment must contain a predetermined overhang that matches the vector. Exact sequences are found in **Supplemental Note, Table 1** (colored in cyan)**.** | NNNN | Optional sequence that can be added if there are not enough bases after the last mutation site to design a reverse primer with an appropriate Tm | NNNN |

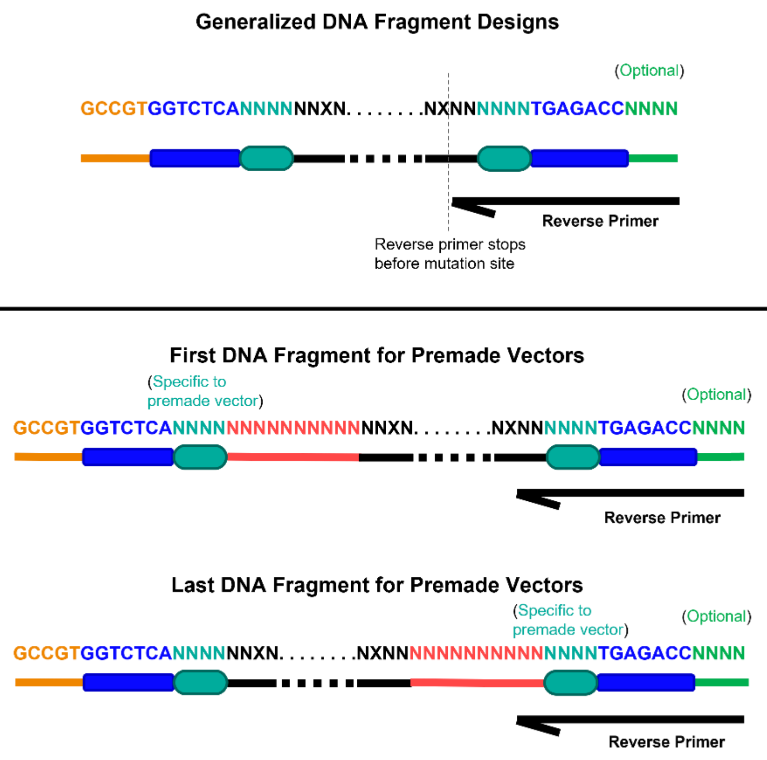


***Supplemental Note, Figure 3.*** *Generalized design schematic of cassettes for Golden Gate-based cassette mutagenesis*

### 2: Cassettes designs for use with pND003, pND004, and pND005

Skip if not using our premade destination vectors.

Cassettes that will be used with pND003, pND004, or pND005 should be designed as described above, with two important exceptions:

First, cassettes should use overhang sequences as outlined in **Supplemental Note, Table 1**. All our destination vectors were made with high-fidelity overhangs from Set 1 from the main text's Table 1 in Potapov *et al*[4]. Once you have chosen a destination vector, the initial overhang from your first cassette and the final overhang from your last cassette should be the ones listed in the sequence for the chosen vector, as seen in **Supplemental Note, Table 1**. The rest of your overhangs should come from the remaining sequences in Set 1, found at the bottom of **Supplemental Note, Table 1**.

Second, your first and last cassettes should include additional sequences, specific to the chosen destination vector, that are shown in red in **Supplemental Note, Table 1.** These code for promoters, terminators, and continuations of tag sequences leading up to the overhang sequences. Note that stop codons do not need to be added to your sequence as these are already coded for in each vector.

| | **First Cassette** | **Last Cassette** |
|---|---|---|
| **pND003** | **GCCGTGGTCTCACGGTAGCGGAGGCGGAGGGTCGGCTAGCCATNNNN......NNNNNNNNTGAGACCNNNN** | **GCCGTGGTCTCANNNNNNNN......NNNNCTCGAGGGGGGCGGATCCGAATGAGACCNNNNN** |
| **pND004** | **GCCGTGGTCTCACGGTCGTCAGACTGTCGATGAAGCCCTGAAAGACGCGCAGACTGGCGGCGGGGGAGGTNNNN.......NNNNNNNNTGAGACCNNNN** | **GCCGTGGTCTCANNNNNNNN......NNNNTGAGATCCGGCTGCTAACAAAGCCCGAAAGGATGAGACCNNNN** |
| **pND005** | **GCCGTGGTCTCATCCTGAAAGNNNN......NNNNNNNNTGAGACCNNNN** | **GCCGTGGTCTCANNNNNNNN......NNNNTGAGTCGTATTACCTCAGCCAAGCTAATTCTGAGACCNNNN** |

| | |
|---|---|
| **High-fidelity overhangs that can be used in conjunction with pND003, pND004, and pND005** | **TGCC, GCAA, ACTA, TTAC, CAGA, TGTG, GAGC, AGGA, ATTC, CGAA, ATAG, AAGG, AACT, AAAA, ACCG** |

***Supplemental Note, Table 1.*** *Sequence specific designs of cassettes for use with pND003, pND004, and pND005. Coloring of bases corresponds to schematics in Supplemental Note, Figure 3.*

**3: Design primers for creation of double stranded cassette DNA**

Next, a single reverse primer must be designed to create double stranded cassette DNA. The primer should not overlap with any mutation site. If mutation sites are located close to the overhang sequences, additional bases might need to be added to the end of a cassette to have a long enough complementary sequence for primer binding (**Supplemental Figure 3**). In this study, we used primers with melting temperatures ($T_m$) ranging from 56°C to 72°C with comparable results. For the RBD library, we used NEB Tm calculator to determine the primer melting temperature and the IDT Oligoanalyzer™ tool to analyze the possible hetero-dimers and the Gibbs free energy of the designed primers. We designed primers to avoid large negative ΔG of any possible mismatch (homodimer, unintended heterodimer, secondary structure) limiting this to > -22 kcal/mol. Additionally, we designed the ΔΔG of the intended heterodimer to any other structure to be <-20 kcal/mol. Inspection of the types of secondary structures may also be helpful to look for sequences which can amplify in an unintended manner from an annealed 3' end (e.g. a hairpin or dimer that anneals the 3' end with an extended 5' end can serve as a template for amplifying products like "primer dimers").

**4. Generate cassette dsDNA from synthetic ssDNA.**

Our experimental protocol below starts at this step, in which dsDNA is generated from single-stranded synthetic DNA fragments. We found that smearing sometimes occurred but that using at least 35 PCR cycles leads to sufficient amplification of the desired product over background. The PCR products are then gel purified and can be stored until the next step.

**Day 1**

1. Dilute the synthetic DNA fragments and the reverse primers to a final concentration of 10 µM in Nuclease Free Water. (See Example 2 for equations used to resuspend the synthetic DNA)

   **Note:** The IDT Resuspension Calculator tool can be used.

2. Generate dsDNA cassettes

   a. Assemble the PCR reactions as follows:

| | |
|---|---|
| Cassette (10 µM) | 1.25 µl |
| Primer reverse (10 µM) | 1.25 µl |
| Q5 2X Master Mix | 12.5 µl |
| Nuclease Free Water | 10 µl |
| **Reaction volume** | **25 µl** |

**Note:** We have used KAPA HiFi Ready Mix (Roche) and Phusion polymerase (NEB) with comparable results.

| **PCR cycling conditions** | | | |
|---|---|---|---|
| **Steps** | **Temperature** | **Time** | **Cycles** |
| Initial Denaturation | 98 °C | 30 s | 1 |
| Denaturation | 98 °C | 10 s | 35 cycles |
| Annealing | 56°C* | 30s | |
| Extension | 72 °C | 1:30 min | |
| Final extension | 72 °C | 2 min | 1 |
| Hold | 4 °C | hold | |

*__Note:__ We have demonstrated generation of dsDNA using primers with a Tm of 56°C as well as 72°C.

b. Purify linear DNA by gel electrophoresis and gel extraction. One clear band is expected for each fragment. Purify all excised fragments using a Nucleo-Spin® Gel and PCR Clean-up kit (Macherey-Nagel). Elute in 20µl Nuclease free water. (See Example 3; if trouble getting the right size band see Troubleshooting 1)

c. Quantify the linear DNA products by measuring $A_{260}$ using a spectrophotometer. We often obtain concentrations of approx. 40 ng/µl.

Section 3: Golden Gate assembly and transformation into *E. Coli*

**Background.** Use a Golden Gate reaction to assemble the dsDNA fragments into your destination vector. This section takes about 5 hours, including bacterial transformation, and can be combined with section 2 for a single day library generation protocol

**Day 1 (Cont.)**

1. Assemble the pieces with a Golden Gate reaction.

a. Per 40 fmol of input DNA, assemble the reaction as follows, adding the enzymes last. (See Example 4 for the calculations used)

| | |
|---|---|
| dsDNA fragments | 40 fmol each |
| Destination vector | 40 fmol |
| 10x T4 DNA ligase buffer | 2.5 µl |
| BsaI-HF-v2 | 1 µl (20 units) |
| T4 DNA ligase | 1 µl (400 units) |
| Nuclease Free Water | up to 25µl |
| **Reaction volume** | **25 µl** |

**Note:** input DNA or total reaction size can be scaled up linearly, as shown in the main text. We primarily scale up just the input DNA when building libraries but .

**Note:** NEBridge Golden Gate Assembly kit (NEB) can also be used with comparable results following manufacturer's instructions.

b. Set the reaction on a thermal cycler using the conditions below:

| **Thermal cycler conditions** | | |
|---|---|---|
| **Temperature** | **Time** | **Cycles** |
| 37 °C | 1 min | 60 cycles |
| 16 °C | 1 min | |
| 37 °C | 5 min | 1 |
| 65 °C | 10 min | 1 |
| 4 °C | infinity and beyond | |

c. To concentrate the DNA, perform a PCR clean-up using a Monarch® PCR & DNA clean-up kit (NEB), eluting in 6 µl of nuclease free water.

2. Transform into *E. coli*.

a. Transform the 6 µl of eluent into electrocompetent cells. (See Example 5)

i. Optionally, transform 0.1 ng of pUC18 (XL1 Blue) or pUC19 (TransforMax) into electrocompetent cells to assess competency.

b. Recover the cells in 1 ml SOC or TB media for 1 h at 37 °C with shaking at ~300 rpm.

c. Do 6 serial dilutions of the recovered cells and plate 10 µl of each dilution on an LB agar plate, containing the appropriate antibiotic, to assess transformation efficiencies. Plate the remainder of the cells on a LB agar square Bioassay dish, containing the appropriate antibiotic, using plating beads. Incubate the plates at 37 °C overnight.

**Day 2**

3. Check the transformation efficiency and if it is above the desired efficiency, scrape the Square Bioassay dish. Add ~3 ml of LB to the plate, using an L shaped cell scraper or a bended glass pasteur pipette, scrape the cells off the agar plate. Collect the cells in a corner and transfer them to a 50ml falcon tube by pipetting. Repeat this process five to seven times for a total 15-20 ml cell volume or until the plate is clear of cells. (See Example 6 to calculate the desired transformation efficiency)
   **Note:** Depending on the destination vector, it might take up to two days to see green colonies. A good control to compare fluorescence of the uncut destination vector is to transform the neat destination vector by itself.

4. Make *E. Coli* glycerol stocks by spinning down 1.5 ml of the scraped cells and resuspending in 700 µl LB media and 300 µl 50 w/v% glycerol.
   **Note:** Since the desired transformation efficiency is 100 fold above the library size, 1.5ml of the scraped cells will have a good representation of the library.

5. Prepare 10 mL of the scraped cells using a ZymoPURE Midiprep kit (Zymo Research) to obtain the plasmid library.

### EXAMPLES

#### Example 1:

In the case of the RBD library, positions 333-416 and 510-541 were not mutated, therefore, they are part of the destination vector. For the RBD we wanted to mutate separately the epitopes of class 1 and class 2 antibodies[7]. Therefore, we divided positions 400 to 509 into 3 cassettes: cassette 1 containing positions 400-436, cassette 2 containing positions 436-472 and cassette 3 containing positions 472-509. This means that positions 400, 436, 472 and 509 can not be mutated (**Figure S2**).

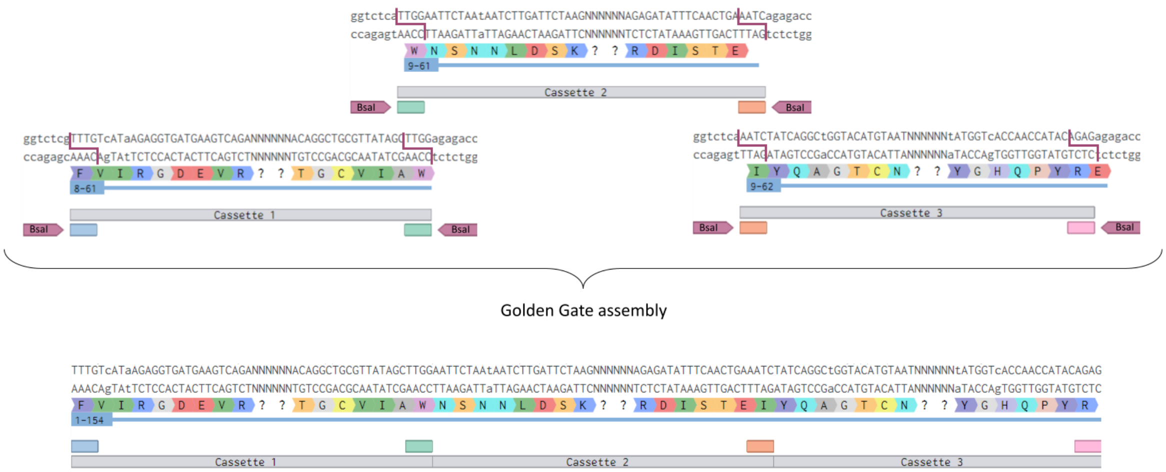


***Supplemental Figure 4.*** *Schematic of cassette assembly and location of the BsaI overhangs. The BsaI overhang at the end of cassette 1 matches the beginning of cassette 2 (green). Since those 4 bp allow the assembly, the codon(s) containing the overhang nucleotides cannot be mutated. Notice that the BsaI sites are located on the outside of each cassette and are not part of the assembled product.*

**Example 2:**

If the ultramer ordered contains 4 nmol of DNA, resuspending in 200 µl will give a 20µM concentration.

$$4\ nmol \times \frac{1\ \mu mol}{10^3\ nmol} \times \frac{l}{20\ \mu mol} \times \frac{10^6 \mu l}{1\ l} = 200 \mu l$$

*The oligo pool ordered contained* 10 pmol of DNA per individual oligo with 1644 oligos. Since we want the final concentration to be per pool:

$$10\ pmol\ \times 1644\ oligos \times \frac{1\ nmol}{10^3\ pmol} =\ 16.44\ nmol$$

Resuspending in 164.4 µl will give a 100µM concentration of the pool.

$$16.44\ nmol \times \frac{1\ \mu mol}{10^3\ nmol} \times \frac{l}{100\ \mu mol} \times \frac{10^6 \mu l}{1\ l} = 164.4\ \mu l$$

**Example 3:**

The fragments for the RBD are ~170 bp, so the 170bp band was excised and gel extracted.

**Example 4:**

*All dilutions are in nuclease free water

| Cassette | Length (bp) | Concentration from PCR (ng/µl) |
|---|---|---|
| 1 | 167 | 33.56 |
| 2 | 168 | 40.24 |
| 3 | 172 | 30.31 |

Cassette 1

$$167\ bp \times \frac{650\ Da}{1\ bp} \times \frac{1\ g}{1\ Da \cdot mol} \times 40 \cdot 10^{-15} mol \times \frac{10^9 ng}{1\ g} = 4.34ng 4.34\ ng \times \frac{1\ \mu l}{33.56ng} = 0.13\ \mu l$$

→ 1.3 µl 1:10 dilution

Cassette 2

$$168\ bp \times \frac{650\ Da}{1\ bp} \times \frac{1\ g}{1\ Da \cdot mol} \times 40 \cdot 10^{-15} mol \times \frac{10^9 ng}{1\ g} = 4.37ng 4.37\ ng \times \frac{1\ \mu l}{40.24ng} = 0.11\ \mu l$$

→ 1.1 µl 1:10 dilution

Cassette 3

$$172\ bp \times \frac{650\ Da}{1\ bp} \times \frac{1\ g}{1\ Da \cdot mol} \times 40 \cdot 10^{-15} mol \times \frac{10^9 ng}{1\ g} = 4.47ng 4.47\ ng \times \frac{1\ \mu l}{40.31\ ng} = 0.11\ \mu l$$

→ 1.1 µl 1:10 dilution

Destination vector

$$7461\ bp \times \frac{650\ Da}{1\ bp} \times \frac{1\ g}{1\ Da \cdot mol} \times 40 \cdot 10^{-15} mol \times \frac{10^9 ng}{1\ g} = 139.98\ ng 139.98\ ng \times \frac{1\ \mu l}{599\ ng} = 0.23\ \mu l$$

→ 2.3 µl 1:10 dilution

| | |
|---|---|
| Cassette 1 | 1.3 µl |
| Cassette 2 | 1.1 µl |
| Cassette 3 | 1.1 µl |
| Destination vector | 2.3 µl |
| T4 DNA ligase buffer | 2.5 µl |
| BsaI-HF-v2 | 1 µl |
| T4 DNA ligase | 1 µl |
| Nuclease Free Water | 14.7 µl |
| **Reaction volume** | **25 µl** |

**Example 5:**

XL1-Blue Competent Cells (Agilent) or TransforMax EC100 Competent Cells (Biosearch Technologies) have been used with comparable results. In our hands when transforming 0.1ng of pUC18 into XL1-Blue we obtain $1.0e^5$ transformants ($1.0e^9$ transformants/µg DNA) and when transforming 0.1ng of pUC19 into TransforMax EC100 we obtain $6.2e^5$ ($6.2e^9$ transformants/µg DNA).

**Example 6:**

The RBD libraries have a theoretical library size on the order of 10k protein-encoding variants. To ensure that all the variants are transformed, we want a 100-fold coverage. Therefore, the minimum desired transformation efficiency is $1e^6$.

### TROUBLESHOOTING

Troubleshooting 1:

If a single band is not observed, increase the extension time and/or increase the number of cycles. If Q5 1X Master Mix (NEB) does not give the desired results, the reaction can be set up with KAPA HiFi Ready Mix (Roche). Alternatively, using "touchdown PCR"[8] may also minimize smearing.

*Supplementary Methods*

### Performance and assessment of Golden Gate reactions

All Golden Gate reactions were performed using a base protocol with some small number of changes which are listed below. The base protocol is as follows. 40 fmol of a destination vector and 40 fmol of each cassette are combined in a PCR tube along with 20 units of BsaI HF-V2, 400 units of T4 DNA ligase, 2.5 µl of 10x T4 ligase buffer, and nuclease-free $H_2O$ up to 25 µl. The reaction mixture is then cycled between a 37 °C and 16 °C PCR step for a total of 60 cycles, followed by a final 5 minute, 37 °C step and a 10 minute, 65 °C step. The resulting DNA is then cleaned and concentrated using an NEB Monarch PCR DNA Cleanup Kit and eluted in 6 µl of nuclease-free $H_2O$. All 6 µl of concentrated DNA are then transformed by electroporation into 50 µl of TransforMax EPI300 *E. coli* cells. Additionally, 0.1 ng of pUC19 in 6 µl of nuclease-free water was transformed into TransforMax cells to assess cell competency. Transformations were carried out in 1 mm electroporation cuvettes (BTX, Cat# 45-0124) with an electroporator (Eppendorf, Cat# 4309000027) set to 1200 V, which generated time constants between 4-6 ms. Immediately after electroporation, cells were resuspended in 1 ml of SOC media (SOB from BD Cat# 244310 with 20mM dextrose), incubated at 37 °C for 1 hour, plated on LB agar containing the appropriate antibiotic, and were incubated overnight at 37 °C. Numbers of transformants and percentages of GFP negative colonies were determined by plating serial dilutions of the transformed cells ranging from $10^2$-$10^8$ fold-diluted and counting the number of green and white colonies in the dilution that had the highest number of colonies totaling between 10 and 100. If no green colonies could be detected, the percentage incorporation was estimated as one over the total number of colonies in the lowest dilution, as extrapolated from the dilution containing all counted colonies. pUC19 transformation efficiencies were similarly determined by counting colonies from serial dilution plating, with the resulting numbers being scaled to reflect the number of transformants per 1 ng of transformed pUC19. For libraries, the remaining transformants not used for dilutions were plated onto bioassay plates and incubated at 37 °C. The next day colonies were scraped from the plates and mini prepped to obtain library DNA. For some experiments, the amount of input DNA or total reaction size was modified from the base protocol; these modifications are noted in the main text. Reaction size and input DNA along with the destination vector and cassettes used in each reaction are listed in **Supplemental Table 1**. The T7 RNAP library was generated with several small additional changes to our base protocol. Two 200 µl reactions (same PCR step reaction; 200 femtomoles of backbone and each insert in each reaction, 20 µl 10x T4 DNA Ligase Buffer, 5 µl BsaI-HFv2, 5 µl T4 DNA Ligase) were pooled, put through a PCR cleanup step, and used in a single electroporation transformation.

### Deep sequencing preparation

Deep sequencing prep was performed as described in the "Method B" protocol from Kowalsky *et al*[9]. In brief, we did two rounds of PCR with an ExoI clean up in between. In the first round we amplify the amplicon using customized primers for each specific sequence. These primers also contain the TruSeq illumina adapters. The second PCR inserts the TruSeq barcodes for sequencing and is performed exactly as described in the “Method B” protocol. The primers used for the deep sequencing preparation  are given in ***Supplementary Table 3***.

The first PCR thermocycler conditions for the RBD library are as follows:

| **PCR cycling conditions** | | | |
|---|---|---|---|
| **Steps** | **Temperature** | **Time** | **Cycles** |
| Initial Denaturation | 98 °C | 60 sec | 1 |
| Denaturation | 98 °C | 10 sec | 18 cycles |
| Annealing | 57°C | 20 sec | |
| Extension | 72 °C | 30 sec | |
| Final extension | 72 °C | 10 min | 1 |
| Hold | 4 °C | hold | |

The first PCR thermocycler conditions for the T7 RNA polymerase library are as follows:

| **PCR cycling conditions** | | | |
|---|---|---|---|
| **Steps** | **Temperature** | **Time** | **Cycles** |
| Initial Denaturation | 98 °C | 60 sec | 1 |
| Denaturation | 98 °C | 10 sec | 16 cycles |
| Annealing | 69°C | 20 sec | |
| Extension | 72 °C | 30 sec | |
| Final extension | 72 °C | 10 min | 1 |

| | | |
|---|---|---|
| Hold | 4 °C | hold |

**DNA deep sequencing analysis**

For the analysis of the RBD library deep sequencing data, sequences were merged using an in-house merging code, essentially as previously described by Haas *et al*[10] The occurrence of mutations at each designed location were counted and summed. From these values, the corresponding distributions were generated (**Figure 3E**).

For analysis of the T7 RNAP library deep sequencing data, sequences were merged using FLASh with a maximum overhang of 100 bp[11]. Merged sequences were filtered to be 416 bp, start with amino acid sequence "EIL" in the appropriate reading frame, contain no "N" basecalls, and to be absent of any premature stop codons. Paired-end reads of length (412-417 bp) were used for comparison to correct lengths as these are the most likely errant lengths, and these showed up at elevated counts relative to outside of this range. The occurrence of mutations at each designed location were counted and summed, including the occurrence of amino acids not in the original design space. From these values, the corresponding distributions were generated (**Figure 4**).

### *Supplementary Table 1*

| Experiment | Rep. | Destination Vector | Insert(s) | Input DNA (fmol) | Size (µl) | Colonies | % GFP Negative | pUC19 Colonies /µg DNA |
|---|---|---|---|---|---|---|---|---|
| Transformants / Cassette Number (Fig. 2A) | 1 | pND003 | PYR1 Cass1+2+3+4 | 40 | 25 | $8.4x10^{5}$ | 99.9 | $3.7x10^{9}$ *(Rep. 1)* $4.5x10^{9}$ *(Rep. 2)* |
| | 2 | | PYR1 Cass1+2+3+4 | | | $8.4x10^{5}$ | 99.9 | |
| | 1 | | PYR1 Cass1+2+3, PYR1 Cass4 | | | $4.9x10^{5}$ | 99.8 | |
| | 2 | | PYR1 Cass1+2+3, PYR1 Cass4 | | | $7.2x10^{5}$ | 99.9 | |
| | 1 | | PYR1 Cass1+2, PYR1 Cass3, | | | $3.4x10^{5}$ | 99.4 | |
| | 2 | | PYR1 Cass1+2, PYR1 Cass3, | | | $6.0x10^{5}$ | 99.9 | |
| | 1 | | PYR1 Cass1, 2, 3, 4 | | | $3.5x10^{5}$ | 99.7 | |
| | 2 | | PYR1 Cass1, 2, 3, 4 | | | 3.9x105 | 99.7 | |
| GG RBD Function Test (Fig. 1D, 1E) | 1 | pIFU037 | RBD WT eBlock (1+2+3) | 40 | 20 | $3x10^{3}$ | 80.0 | NA |
| | 2 | | | | | $1x10^{4}$ | 91.6 | |
| | 3 | | | | | $1.1x10^{6}$ | 96.0 | |
| | 4 | | | | | $5.3x10^{5}$ | 98.0 | |
| Transformants / Reaction Size (Fig. 2B) | 1 | pND003 | PYR1 Cass1+2+3+4 | 40 | 25 | $3.2x10^{5}$ | 96.9 | $8.3x10^{9}$ *(Rep. 1)* $4.9x10^{9}$ *(Rep. 2)* $7.1x10^{9}$ *(Rep. 3)* |
| | 2 | pND003 | PYR1 Cass1+2+3+4 | 40 | 25 | $9x10^{5}$ | 98.9 | |
| | 3 | pND003 | PYR1 Cass1+2+3+4 | 40 | 25 | $9x10^{5}$ | 98.9 | |
| | 1 | pND003 | PYR1 Cass1+2+3+4 | 160 | 100 | $2x10^{6}$ | 99.5 | |
| | 2 | pND003 | PYR1 Cass1+2+3+4 | 160 | 100 | $2.4x10^{6}$ | 99.6 | |
| | 3 | pND003 | PYR1 Cass1+2+3+4 | 160 | 100 | $2.7x10^{6}$ | 99.6 | |
| | 1 | pND003 | PYR1 Cass1+2+3+4 | 320 | 200 | $8.2x10^{6}$ | 99.9 | |
| | 2 | pND003 | PYR1 Cass1+2+3+4 | 320 | 200 | $2.3x10^{7}$ | 99.9 | |
| | 3 | pND003 | PYR1 Cass1+2+3+4 | 320 | 200 | $1.9x10^{7}$ | 99.9 | |
| Transformants / Input DNA (Fig. 2C) | 1 | pND003 | PYR1 Cass1+2+3+4 | 80 | 25 | $7x10^{5}$ | 98.6 | |
| | 2 | pND003 | PYR1 Cass1+2+3+4 | 80 | 25 | $2.9x10^{6}$ | 99.7 | |
| | 3 | pND003 | PYR1 Cass1+2+3+4 | 80 | 25 | $4x10^{6}$ | 99.8 | |
| | 1 | pND003 | PYR1 Cass1+2+3+4 | 160 | 25 | $5.8x10^{6}$ | 99.7 | |
| | 2 | pND003 | PYR1 Cass1+2+3+4 | 160 | 25 | $2.7x10^{7}$ | 99.8 | |
| | 3 | pND003 | PYR1 Cass1+2+3+4 | 160 | 25 | $6.2x10^{6}$ | 99.8 | |
| RBD Library (Fig. 3A) | 1 | pIFU037 | RBD ultramer 1+2+3 | 200 | 25 | $5.6x10^{6}$ | 99.9 | NA |
| | 1 | | RBD ultramer 1+2+3 | | | $2x10^{6}$ | 99.9 | |
| | 1 | | RBD oligo pool 1+2+3 | 400 | 50 | $1.6x10^{7}$ | 98.0 | |
| | 2 | | | | | $2.3x10^{7}$ | 92.0 | |
| eBlock, Ultramer, oligo pool comparison (Fig. 3D) | 1 | pIFU037 | WT RBD eBlock 1+2+3 | 40 | 25 | $1.5x10^{7}$ | 99.9 | $6.2x10^{9}$ *(Rep. 1)* $5x10^{9}$ *(Rep. 2)* $2.5x10^{9}$ *(Rep. 3)* |
| | 2 | | | | | $2.5x10^{6}$ | 99.9 | |
| | 3 | | | | | $3.4x10^{6}$ | 99.9 | |
| | 1 | | WT RBD Ultramer 1+2+3 | | | $3.4x10^{6}$ | 99.9 | |
| | 2 | | | | | $1.5x10^{6}$ | 99.9 | |
| | 3 | | | | | $1.4x10^{6}$ | 99.9 | |
| | 1 | | RBD oligo pool 1+2+3 | | | $3.0x10^{6}$ | 99.9 | |
| | 2 | | | | | $1.1x10^{6}$ | 99.9 | |
| | 3 | | | | | $2.2x10^{6}$ | 99.9 | |
| T7 Library (Fig. 4A) | 1 | pZB258 | T7 Cass 1 + Cass 2 | 400 | 200 | $5.6x10^{7}$ | 99.6 | NA |

*Supplementary Table 2*

| | **Library 1** | **Library 2** | **Library 3** |
|---|---|---|---|
| **DNA type** | Ultramer | Ultramer | Oligo pool |
| **Number of designed mutations** | 7776 | 20736 | 17725 |
| **Number of transformants obtained** | $5.6x10^6$ | $2x10^6$ | $2x10^7$ |
| **Cassette incorporation efficiency** | 99.9% | 99.9% | 99.9% |
| **Library coverage** | 100% (7776/7776) | 96.7% (20043/20736) | 99.9% (17701/17725) |
| **% desired reads** (designed variants - right mutation and number of expected mutations) | 86.8% (824718/949555) | 83.9% (594753/708766) | 80.9% (2607572/3224158) |
| **% chimera reads** (variants with the correct mutations but with more than 3 mutations in cassette 3) | 0% (0/949555) | 0% (0/708766) | 8.9% (289648/3224158) |
| **% WT sequence reads** | 0.22% (1798/824718) | 0.06% (347/594753) | 0.02% (483/2607572) |

### *Supplementary Table 3*

| Primer Name | Sequence |
|---|---|
| NDD-0126 pJS755 Backbone HA Rev | ACGTTTCACTTTCGGTCTCAACCGCTGGCGGCG |
| NDD-0125 pJS755 Backbone HA Fwd | TTCTGCGTTTATAGGTCTCAAGGAAGCTGAGTTGGCTGC |
| NDD-0124 pJS637 Backbone HA Rev | ACGTTTCACTTTCGGTCTCAACCGCCTCCACCAGAG |
| NDD-0123 pJS637 Backbone HA Fwd | TTCTGCGTTTATAGGTCTCACGAACAAAAGCTTATTTCTGAAGAGG |
| NDD-0122 pACL032 Backbone HA Rev | ACGTTTCACTTTCGGTCTCAAGGAGGCTTGCTTCAAGCT |
| NDD-0121 pACL032 Backbone HA Fwd | TTCTGCGTTTATAGGTCTCAATTCCGGGCGAATTTCTTATGATT |
| NDD-0120 GFP HA Rev | TGAGACCTATAAACGCAGAAAGG |
| NDD-0119 GFP HA Fwd | TGAGACCGAAAGTGAAACGT |
| NDD-0116 pJS755 BSa1 Removal Rev | CTTACCGACTTCAGCGAGTCCGTTATAGCCTTTATC |
| NDD-0115 pJS755 BSa1 Removal Fwd | GATAAAGGCTATAACGGACTCGCTGAAGTCGGTAAG |
| ZTB Cass1_for | GCATCGGGTCTCATTCGCAAGCGTTGCGCTGBNCATTGGGTAACTCCT GATGGTTTCCCTGBNTGGNDTGAATACAAGAAGCCTATTCAGACGCG CTTGAACCTGATGTTCCTCGGTCAGTTCCGCTTACAGCCTACCATTAAC ACCAACAAAGATAGCGAGAT |
| ZTB Cass1_Q737_for | GCATCGGGTCTCATTCGCAAGCGTTGCGCTGBNCATTGGGTAACTCCT GATGGTTTCCCTGBNTGGCAAGAATACAAGAAGCCTATTCAGACGCG CTTGAACCTGATGTTCCTCGGTCAGTTCCGCTTACAGCCTACCATTAAC ACCAACAAAGATAGCGAGAT |
| ZTB Cass1_rev | GCTACGGGTCTCAATCTCGCTATCTTTGTTGGTGTTAATGGTAGGCTG TAAGCGG |
| ZTB Cass2_for | GCATCGGGTCTCAAGATTGATGCACACAAACAGGAGTCTGGTVTAGC TCCTMRNNDTGTACACRSYVRKGACGGTAGCCACCTTCGTAAGACTG TAGTGTGGGCACACGAGAAGTACGGAATCGAATCTTTTGCACTGATT CACGACTCCTTCGGTACCATTCCGGCTGACGCTGCGAACCTGTTCAAA GCAGTGCGC |
| ZTB Cass2_rev | GCTACGGGTCTCAAGCGATCTGGTCGTAAHNATCAGCCAGTACATCA CAAGACTCATATGTGTCAACCATAGTTTCGCGCACTGCTTTGAACAGG TTCGCAGCGTCAGCCGGAATGGTACCGAAGGA |
| ZTB NGS_for | GTTCAGAGTTCTACAGTCCGACGATCGAGAGATTCTTCGCAAGCGTTG CG |
| ZTB NGS_rev | CCTTGGCACCCGAGAATTCCAGTGCAACTGGTCAGCGATCTGG |
| IFU-159 RBD cass1_rev | ACAGGCTGCGTTATAGCTTGGAGAGACCATAGATATGATGTAGCGTA GC |
| IFU-160 RBD cass2_rev | TTCGAGAGAGATATTTCAACTGAAATCAGAGACCATGCTAGAGCGTA TGATGTAGCGTAGC |
| IFU-161 RBD cass3_rev | CAACCATACAGAGAGAGACCATGCTAGAGCGTATGATGTAGCGTAGC |
| IFU-162 RBD_fwd | GCATCGTTCCCTACTGCTCGTACATGC |
| IFU-163 RBD NGS_fwd | GTTCAGAGTTCTACAGTCCGACGATCTAGAGGTGATGAAGTCAGA |
| IFU-164 RBD NGS_rev | CCTTGGCACCCGAGAATTCCATACTACTACTCTGTATGGTTG |
| Forward Outer Primer | AATGATACGGCGACCACCGAGATCTACACGTTCAGAGTTCTACAGTCC GACGATC |
| Reverse Outer Primer | GGCATACGAGATNNNNNNGTGACTGGAGTTCCTTGGCACCCGAGAA TTCCA ‘‘NNNNNN’’ is the barcode |

***Supplementary Data***

**Cassettes**

| |
|---|
| PYR1 Cass1<br>TTCCCTACTGCTCGTACATCCGGTCTCACGGTAGCGGAGGCGGAGGGTCGGCTAGCCATATGCCTTC<br>GGAGTTAACACCAGAAGAACGATCGGAACTAAAAAACTCAATCGCCGAGTTCCACACATACCAACT<br>CGATCCAGGAAGCTGTTCATCACTCCACGCGCAACGAATCCACGCGCCTCCGGAACTCGTCTGTGAG<br>ACCATAGCGGTTCTCACCCCTCAACACCTGCGCTGTCCGCACCGTTTGAAGTAAATTAGTTGAGGAT<br>TTAGCAGTGCTATCATGTGATCTCCAAATTAAAACATACCGTTCCATGAGGGCTAGAATTACTTACC<br>GGCCTTCACCATGCCTGTACTATACGAACCCACTCTC |
| PYR1 Cass2<br>TTCCCTACTGCTCGTACATCCGGTCTCATCTGGTCAATCGTACGACGATTCGACAAACCACAAACAT<br>ACAAACCGTTCATCAAATCCTGCTCCGTCGAACAAAACTTCGAGATGCGCGTCGGATGCACGCGCG<br>ACGTGATCGTCATCAGTGGATTACCGGCGTCTACATCAACGGAAAGACTCGATATACTCGACGACG<br>AACGGAGAGTTACTGAGACCATAGCGGTTCTCACCCCTCAACACCTGCGCTGTCCGCACCGTTTGAA<br>GTAAATTAGTTGAGGATTTAGCAGTGCTATCATGTGATCTCCAAATTAAAACATACCGTTCCATGAG<br>GGCTAGAATTACTTACCGGCCTTCACCATGCCTGTACTATACGAACCCACTCTC |
| PYR1 Cass3<br>TTCCCTACTGCTCGTACATCCGGTCTCATTACCGGATTCAGTATCATCGGAGGCGAACATAGGCTGA<br>CGAATTACAAATCCGTTACGACGGTGCATCGGTTCGAGAAAGAGAATCGGATCTGGACGGTGGTTT<br>TGGAATCTTACGTCGTTGATATGCCTGAGACCATAGCGGTTCTCACCCCTCAACACCTGCGCTGTCC<br>GCACCGTTTGAAGTAAATTAGTTGAGGATTTAGCAGTGCTATCATGTGATCTCCAAATTAAAACATA<br>CCGTTCCATGAGGGCTAGAATTACTTACCGGCCTTCACCATGCCTGTACTATACGAACCCACTCTC |
| PYR1 Cass4<br>TTCCCTACTGCTCGTACATCCGGTCTCATGCCGGAAGGTAACTCGGAGGATGATACTCGTATGTTTG<br>CTGATACGGTTGTGAAGCTTAATTTGCAGAAACTCGCGACGGTTGCTGAAGCTATGGCTCGTAACTC<br>CGGTGACGGAAGTGGTTCTCAGGTGACGCTCGAGGGGGGCGGATCCGAATGAGACCATAGCGGTTC<br>TCACCCCTCAACACCTGCGCTGTCCGCACCGTTTGAAGTAAATTAGTTGAGGATTTAGCAGTGCTAT<br>CATGTGATCTCCAAATTAAAACATACCGTTCCATGAGGGCTAGAATTACTTACCGGCCTTCACCATG<br>CCTGTACTATACGAACCCACTCTC |
| PYR1 Cass1+2<br>TTCCCTACTGCTCGTACATCCGGTCTCACGGTAGCGGAGGCGGAGGGTCGGCTAGCCATATGCCTTC<br>GGAGTTAACACCAGAAGAACGATCGGAACTAAAAAACTCAATCGCCGAGTTCCACACATACCAACT<br>CGATCCAGGAAGCTGTTCATCACTCCACGCGCAACGAATCCACGCGCCTCCGGAACTCGTCTGGTCA<br>ATCGTACGACGATTCGACAAACCACAAACATACAAACCGTTCATCAAATCCTGCTCCGTCGAACAA<br>AACTTCGAGATGCGCGTCGGATGCACGCGCGACGTGATCGTCATCAGTGGATTACCGGCGTCTACAT<br>CAACGGAAAGACTCGATATACTCGACGACGAACGGAGAGTTACTGAGACCATAGCGGTTCTCACCC<br>CTCAA |
| PYR1 Cass1+2+3<br>TTCCCTACTGCTCGTACATCCGGTCTCACGGTAGCGGAGGCGGAGGGTCGGCTAGCCATATGCCTTC<br>GGAGTTAACACCAGAAGAACGATCGGAACTAAAAAACTCAATCGCCGAGTTCCACACATACCAACT<br>CGATCCAGGAAGCTGTTCATCACTCCACGCGCAACGAATCCACGCGCCTCCGGAACTCGTCTGGTCA<br>ATCGTACGACGATTCGACAAACCACAAACATACAAACCGTTCATCAAATCCTGCTCCGTCGAACAA<br>AACTTCGAGATGCGCGTCGGATGCACGCGCGACGTGATCGTCATCAGTGGATTACCGGCGTCTACAT<br>CAACGGAAAGACTCGATATACTCGACGACGAACGGAGAGTTACCGGATTCAGTATCATCGGAGGCG<br>AACATAGGCTGACGAATTACAAATCCGTTACGACGGTGCATCGGTTCGAGAAAGAGAATCGGATCT<br>GGACGGTGGTTTTGGAATCTTACGTCGTTGATATGCCTGAGACCATAGCGGTTCTCACCCCTCAA |

| |
|---|
| PYR1 Cass1+2+3+4<br>TTCCCTACTGCTCGTACATCCGGTCTCACGGTAGCGGAGGCGGAGGGTCGGCTAGCCATATGCCTTC<br>GGAGTTAACACCAGAAGAACGATCGGAACTAAAAAACTCAATCGCCGAGTTCCACACATACCAACT<br>CGATCCAGGAAGCTGTTCATCACTCCACGCGCAACGAATCCACGCGCCTCCGGAACTCGTCTGGTCA<br>ATCGTACGACGATTCGACAAACCACAAACATACAAACCGTTCATCAAATCCTGCTCCGTCGAACAA<br>AACTTCGAGATGCGCGTCGGATGCACGCGCGACGTGATCGTCATCAGTGGATTACCGGCGTCTACAT<br>CAACGGAAAGACTCGATATACTCGACGACGAACGGAGAGTTACCGGATTCAGTATCATCGGAGGCG<br>AACATAGGCTGACGAATTACAAATCCGTTACGACGGTGCATCGGTTCGAGAAAGAGAATCGGATCT<br>GGACGGTGGTTTTGGAATCTTACGTCGTTGATATGCCGGAAGGTAACTCGGAGGATGATACTCGTAT<br>GTTTGCTGATACGGTTGTGAAGCTTAATTTGCAGAAACTCGCGACGGTTGCTGAAGCTATGGCTCGT<br>AACTCCGGTGACGGAAGTGGTTCTCAGGTGACGCTCGAGGGGGGCGGATCCGAATGAGACCATAGC<br>GGTTCTCACCCCTCAA |
| RBD eblock Cass1<br>TTCCCTACTGCTCGTACATCCGGTCTCATTTGTAATTAGAGGTGATGAAGTCAGACAAATCGCTCCA<br>GGGCAAACTGGAAAGATTGCTGATTATAATTATAAATTACCAGATGATTTTACAGGCTGCGTTATAG<br>CTTGGAGAGACCATAGCGGTTCTCACCCCTCAA |
| RBD eblock Cass2<br>TTCCCTACTGCTCGTACATCCGGTCTCATTGGAATTCTAACAATCTTGATTCTAAGGTTGGTGGTAAT<br>TATAATTACCTGTATAGATTGTTTAGGAAGTCTAATCTCAAACCTTTTGAGAGAGATATTTCAACTG<br>AAATCAGAGACCATAGCGGTTCTCACCCCTCAA |
| RBD eblock Cass3<br>TTCCCTACTGCTCGTACATCCGGTCTCAAATCTATCAGGCCGGTAACACACCTTGTAATGGTGTTGCA<br>GGTTTTAATTGTTACTTTCCTTTACAATCATATGGTTTCCGACCCACTTATGGTGTTGGTCACCAACC<br>ATACAGAGAGAGACCATAGCGGTTCTCACCCCTCAA |
| RBD Ultramer Cass1<br>GCATCGGGTCTCGTTTGTCATAAGAGGTGATGAAGTCAGACAAATCGCTCCAGGGCAAACTGGAAA<br>GATTGCTGACTACAATTACAAGTTACCAGATGATTTTACAGGCTGCGTTATAGCTTGGAGAGACCAT<br>AGATATGATGTA |
| RBD Ultramer Cass2<br>GCATCGGGTCTCATTGGAATTCTAATAATCTTGATTCTAAGGTAGGTGGAAATTACAATTACCTGTA<br>TAGATTGTTTAGGAAGTCTAATCTCAAACCTTTCGAGAGAGATATTTCAACTGAAATCAGAGACCAT<br>GCTAGAGCGTATGATGTA |
| RBD opool Cass1<br>GCATCGTTCCCTACTGCTCGTACATGCGGTCTCGTTTGTCATAAGAGGTGATGAAGTCAGACAAATC<br>GCTCCAGGGCAAACTGGAAAGATTGCTGACTACAATTACAAGTTACCAGATGATTTTACAGGCTGC<br>GTTATAGCTTGGAGAGACCATAGATATGATGTA |
| RBD opool Cass2<br>GCATCGTTCCCTACTGCTCGTACATGCGGTCTCATTGGAATTCTAATAATCTTGATTCTAAGGTAGGT<br>GGAAATTACAATTACCTGTATAGATTGTTTAGGAAGTCTAATCTCAAACCTTTCGAGAGAGATATTT<br>CAACTGAAATCAGAGACCATGCTAGAGCGTATGATGTA |
| RBD opool Cass3<br>GCATCGTTCCCTACTGCTCGTACATGCGGTCTCAAATCTATCAGGCTGGTAACACACCTTGTAATGG<br>TGTTGCAGGTTTTAATTGTTACTTTCCTTTACAATCATATGGCTTCCGACCCACTTATGGTGTAGGTC<br>ACCAACCATACAGAGAGAGACCATGCTAGAGCGTATGATGTA |

T7 RNAP Cass1
GCATCGGGTCTCATTCGCAAGCGTTGCGCTGBNCATTGGGTAACTCCTGATGGTTTCCCTGBNTGGN
DTGAATACAAGAAGCCTATTCAGACGCGCTTGAACCTGATGTTCCTCGGTCAGTTCCGCTTACAGCC
TACCATTAACACCAACAAAGATAGCGAGATTGAGACCCGTAGC

T7 RNAP Cass2
GCATCGGGTCTCAAGATTGATGCACACAAACAGGAGTCTGGTVTAGCTCCTMRNNDTGTACACRSY
VRKGACGGTAGCCACCTTCGTAAGACTGTAGTGTGGCACACGAGAAGTACGGAATCGAATCTTTT
GCACTGATTCACGACTCCTTCGGTACCATTCCGGCTGACGCTGCGAACCTGTTCAAAGCAGTGCGCG
AAACTATGGTTGACACATATGAGTCTTGTGATGTACTGGCTGATNDTTACGACCAGATCGCTTGAGA
CCCGTAGC

**Destination Vectors**

pND003
TAATTATTTTTATAGCACGTGATGAAAAGGACCCAGGTGGCACTTTTCGGGGAAATGTGCGCGGAAC
CCCTATTTGTTTATTTTTCTAAATACATTCAAATATGTATCCGCTCATGAGACAATAACCCTGATAAA
TGCTTCAATAATATTGAAAAAGGAAGAGTATGAGCCATATTCAACGGGAAACGTCTTGCTCTAGGC
CGCGATTAAATTCCAACATGGATGCTGATTTATATGGGTATAAATGGGCTCGCGATAATGTCGGGCA
ATCAGGTGCGACAATCTATCGATTGTATGGGAAGCCCGATGCGCCAGAGTTGTTTCTGAAACATGGC
AAAGGTAGCGTTGCCAATGATGTTACAGATGAGATGGTCAGACTAAACTGGCTGACGGAATTTATG
CCTCTTCCGACCATCAAGCATTTTATCCGTACTCCTGATGATGCATGGTTACTCACCACTGCGATCCC
CGGGAAAACAGCATTCCAGGTATTAGAAGAATATCCTGATTCAGGTGAAAATATTGTTGATGCGCT
GGCAGTGTTCCTGCGCCGGTTGCATTCGATTCCTGTTTGTAATTGTCCTTTTAACAGCGATCGCGTAT
TTCGTCTCGCTCAGGCGCAATCACGAATGAATAACGGTTTGGTTGATGCGAGTGATTTTGATGACGA
GCGTAATGGCTGGCCTGTTGAACAAGTCTGGAAAGAAATGCATAAACTTTTGCCATTCTCACCGGAT
TCAGTCGTCACTCATGGTGATTTCTCACTTGATAACCTTATTTTTGACGAGGGGAAATTAATAGGTTG
TATTGATGTTGGACGAGTCGGAATCGCAGACCGATACCAGGATCTTGCCATCCTATGGAACTGCCTC
GGTGAGTTTTCTCCTTCATTACAGAAACGGCTTTTTCAAAAATATGGTATTGATAATCCTGATATGAA
TAAATTGCAGTTTCATTTGATGCTCGATGAGTTTTTCTAACCTCAGCAACTGTCAACCAAGTTTACTC
ATATATACTTTAGATTGATTTAAAACTTCATTTTTAATTTAAAAGGATCTAGGTGAAGATCCTTTTTG
ATAATCTCATGACCAAAATCCCTTAACGTGAGTTTTCGTTCCACTGAGCGTCAGACCCCGTAGAAAA
GATCAAAGGATCTTCTTGAGATCCTTTTTTTCTGCGCGTAATCTGCTGCTTGCAAACAAAAAAACCA
CCGCTACCAGCGGTGGTTTGTTTGCCGGATCAAGAGCTACCAACTCTTTTTCCGAAGGTAACTGGCT
TCAGCAGAGCGCAGATACCAAATACTGTTCTTCTAGTGTAGCCGTAGTTAGGCCACCACTTCAAGAA
CTCTGTAGCACCGCCTACATACCTCGCTCTGCTAATCCTGTTACCAGTGGCTGCTGCCAGTGGCGAT
AAGTCGTGTCTTACCGGGTTGGACTCAAGACGATAGTTACCGGATAAGGCGCAGCGGTCGGGCTGA
ACGGGGGGTTCGTGCACACAGCCCAGCTTGGAGCGAACGACCTACACCGAACTGAGATACCTACAG
CGTGAGCTATGAGAAAGCGCCACGCTTCCCGAAGGGAGAAAGGCGGACAGGTATCCGGTAAGCGG
CAGGGTCGGAACAGGAGAGCGCACGAGGGAGCTTCCAGGGGGAAACGCCTGGTATCTTTATAGTCC
TGTCGGGTTTCGCCACCTCTGACTTGAGCGTCGATTTTTGTGATGCTCGTCAGGGGGGCGGAGCCTA
TGGAAAAACGCCAGCAACGCGGCCTTTTTACGGTTCCTGGCCTTTTGCTGGCCTTTTGCTCACATGTT
CTTTCCTGCGTTATCCCCTGATTCTGTGGATAACCGTATTACCGCCTTTGAGTGAGCTGATACCGCTC
GCCGCAGCCGAACGACCGAGCGCAGCGAGTCAGTGAGCGAGGAAGCGGAAGAGCGCCCAATACGC
AAACCGCCTCTCCCCGCGCGTTGGCCGATTCATTAATGCAGCTGGCACGACAGGTTTCCCGACTGGA
AAGCGGGCAGTGAGCGCAACGCAATTAATGTGAGTTAGCTCACTCATTAGGCACCCCAGGCTTTAC
ACTTTATGCTTCCGGCTCGTATGTTGTGTGGAATTGTGAGCGGATAACAATTTCACACAGGAAACAG
CTATGACCATGATTACGCCAAGCTCGAAATTAACCCTCACTAAAGGGAACAAAAGCTGGTACCAAT
TCCTTGAATTTTCAAAAATTCTTACTTTTTTTTTGGATGGACGCAAAGAAGTTTAATAATCATATTAC
ATGGCATTACCACCATATACATATCCATATCTAATCTTACTTATATGTTGTGGAAATGTAAAGAGCC
CCATTATCTTAGCCTAAAAAAACCTTCTCTTTGGAACTTTCAGTAATACGCTTAACTGCTCATTGCTA
TATTGAAGTACGGATTAGAAGCCGCCGAGCGGGTGACAGCCCTCCGAAGGAAGACTCTCCTCCGTG

CGTCCTCGTCTTCACCGGTCGCGTTCCTGAAACGCAGATGTGCCTCGCGCCGCACTGCTCCGAACAA
TAAAGATTCTACAATACTAGCTTTTATGGTTATGAAGAGGAAAAATTGGCAGTAACCTGGCCCCACA
AACCTTCAAATGAACGAATCAAATTAACAACCATAGGATGATAATGCGATTAGTTTTTTAGCCTTAT
TTCTGGGGTAATTAATCAGCGAAGCGATGATTTTTGATCTATTAACAGATATATAAATGCAAAAACT
GCATAACCACTTTAACTAATACTTTCAACATTTTCGGTTTGTATTACTTCTTATTCAAATGTAATAAA
AGTATCAACAAAAAATTGTTAATATACCTCTATACTTTAACGTCAAGGAGAAAAAACCCCGGATCG
AATTCCCTACTTCATACATTTTCAATTAAGATGCAGTTACTTCGCTGTTTTTCAATATTTTCTGTTATT
GCTTCAGTTTTAGCACAGGAACTGACAACTATATGCGAGCAAATCCCCTCACCAACTTTAGAATCGA
CGCCGTACTCTTTGTCAACGACTACTATTTTGGCCAACGGGAAGGCAATGCAAGGAGTTTTTGAATA
TTACAAATCAGTAACGTTTGTCAGTAATTGCGGTTCTCACCCCTCAACAACTAGCAAAGGCAGCCCC
ATAAACACACAGTATGTTTTTAAGGACAATAGCTCGACGATTGAAGGTAGATACCCATACGACGTTC
CAGACTACGCTCTGCAGGCTAGTGGTGGAGGAGGCTCTGGTGGAGGCGGTTGAGACCGAAAGTGAA
ACGTGATTTCATGCGTCATTTTGAACATTTTGTAAATCTTATTTAATAATGTGTGCGGCAATTCACAT
TTAATTTATGAATGTTTTCTTAACATCGCGGCAACTCAAGAAACGGCAGGTTCGGATCTTAGCTACT
AGAGAAAGAGGAGAAATACTAGATGCGTAAAGGCGAAGAGCTGTTCACTGGTGTCGTCCCTATTCT
GGTGGAACTGGATGGTGATGTCAACGGTCATAAGTTTTCCGTGCGTGGCGAGGGTGAAGGTGACGC
AACTAATGGTAAACTGACGCTGAAGTTCATCTGTACTACTGGTAAACTGCCGGTTCCTTGGCCGACT
CTGGTAACGACGCTGACTTATGGTGTTCAGTGCTTTGCTCGTTATCCGGACCATATGAAGCAGCATG
ACTTCTTCAAGTCCGCCATGCCGGAAGGCTATGTGCAGGAACGCACGATTTCCTTTAAGGATGACGG
CACGTACAAAACGCGTGCGGAAGTGAAATTTGAAGGCGATACCCTGGTAAACCGCATTGAGCTGAA
AGGCATTGACTTTAAAGAGGACGGCAATATCCTGGGCCATAAGCTGGAATACAATTTTAACAGCCA
CAATGTTTACATCACCGCCGATAAACAAAAAAATGGCATTAAAGCGAATTTTAAAATTCGCCACAA
CGTGGAGGATGGCAGCGTGCAGCTGGCTGATCACTACCAGCAAAACACTCCAATCGGTGATGGTCC
TGTTCTGCTGCCAGACAATCACTATCTGAGCACGCAAAGCGTTCTGTCTAAAGATCCGAACGAGAAA
CGCGATCATATGGTTCTGCTGGAGTTCGTAACCGCAGCGGGCATCACGCATGGTATGGATGAACTGT
ACAAATGACCAGGCATCAAATAAAACGAAAGGCTCAGTCGAAAGACTGGGCCTTTCGTTTTATCTG
TTGTTTGTCGGTGAACGCTCTCTACTAGAGTCACACTGGCTCACCTTCGGGTGGGCCTTTCTGCGTTT
ATAGGTCTCACGAACAAAAGCTTATTTCTGAAGAGGACTTGTAATAGAGATCTGATAACAACAGTG
TAGATGTAACAAAATCGACTTTGTTCCCACTGTACTTTTAGCTCGTACAAAATACAATATACTTTTCA
TTTCTCCGTAAACAACATGTTTTCCCATGTAATATCCTTTTCTATTTTTCGTTCCGTTACCAACTTTAC
ACATACTTTATATAGCTATTCACTTCTATACACTAAAAAACTAAGACAATTTTAATTTTGCTGCCTGC
CATATTTCAATTTGTTATAAATTCCTATAATTTATCCTATTAGTAGCTAAAAAAAGATGAATGTGAAT
CGAATCCTAAGAGAATTGAGCTCCAATTCGCCCTATAGTGAGTCGTATTACAATTCACTGGCCGTCG
TTTTACAACGTCGTGACTGGGAAAACCCTGGCGTTACCCAACTTAATCGCCTTGCAGCACATCCCCC
TTTCGCCAGCTGGCGTAATAGCGAAGAGGCCCGCACCGATCGCCTTTCCCAACAGTTGCGCAGCCTG
AATGGCGAATGGACGCGCCCTGTAGCGGCGCATTAAGCGCGGCGGGTGTGGTGGTTACGCGCAGCG
TGACCGCTACACTTGCCAGCGCCCTAGCGCCCGCTCCTTTCGCTTTCTTCCCTTCCTTTCTCGCCACGT
TCGCCGGCTTTCCCCGTCAAGCTCTAAATCGGGGGCTCCCTTTAGGGTTCCGATTTAGTGCTTTACGG
CACCTCGACCCCAAAAAACTTGATTAGGGTGATGGTTCACGTAGTGGGCCATCGCCCTGATAGACG
GTTTTTCGCCCTTTGACGTTGGAGTCCACGTTCTTTAATAGTGGACTCTTGTTCCAAACTGGAACAAC
ACTCAACCCTATCTCGGTCTATTCTTTTGATTTATAAGGGATTTTGCCGATTTCGGCCTATTGGTTAA
AAAATGAGCTGATTTAACAAAAATTTAACGCGAATTTTAACAAAATATTAACGCTTACAATTTCCTG
ATGCGGTATTTTCTCCTTACGCATCTGTGCGGTATTTCACACCGCATAGATCGGCAAGTGCACAAAC
AATACTTAAATAAATACTACTCAGTAATAACCTATTTCTTAGCATTTTTGACGAAATTTGCTATTTTG
TTAGAGTCTTTTACACCATTTGTCTCCACACCTCCGCTTACATCAACACCAATAACGCCATTTAATCT
AAGCGCATCACCAACATTTTCTGGCGTCAGTCCACCAGCTAACATAAAATGTAAGCTTTCGGGGCTC
TCTTGCCTTCCAACCCAGTCAGAAATCGAGTTCCAATCCAAAAGTTCACCTGTCCCACCTGCTTCTGA
ATCAAACAAGGGAATAAACGAATGAGGTTTCTGTGAAGCTGCACTGAGTAGTATGTTGCAGTCTTTT
GGAAATACGAGTCTTTTAATAACTGGCAAACCGAGGAACTCTTGGTATTCTTGCCACGACTCATCTC
CATGCAGTTGGACGATATCAATGCCGTAATCATTGACCAGAGCCAAAACATCCTCCTTAGGTTGATT
ACGAAACACGCCAACCAAGTATTTCGGAGTGCCTGAACTATTTTTATATGCTTTTACAAGACTTGAA
ATTTTCCTTGCAATAACCGGGTCAATTGTTCTCTTTCTATTGGGCACACATATAATACCCAGCAAGTC
AGCATCGGAATCTAGAGCACATTCTGCGGCCTCTGTGCTCTGCAAGCCGCAAACTTTCACCAATGGA
CCAGAACTACCTGTGAAATTAATAACAGACATACTCCAAGCTGCCTTTGTGTGCTTAATCACGTATA
CTCACGTGCTCAATAGTCACCAATGCCCTCCCTCTTGGCCCTCTCCTTTTCTTTTTCGACCGAATTAA

TTCTTAATCGGCAAAAAAAGAAAAGCTCCGGATCAAGATTGTACGTAAGGTGACAAGCTATTTTTCA
ATAAAGAATATCTTCCACTACTGCCATCTGGCGTCATAACTGCAAAGTACACATATATTACGATGCT
GTTCTATTAAATGCTTCCTATATTATATATATAGTAATGTCGTGATCTATGGTGCACTCTCAGTACAA
TCTGCTCTGATGCCGCATAGTTAAGCCAGCCCCGACACCCGCCAACACCCGCTGACGCGCCCTGACG
GGCTTGTCTGCTCCCGGCATCCGCTTACAGACAAGCTGTGACCGTCTCCGGGAGCTGCATGTGTCAG
AGGTTTTCACCGTCATCACCGAAACGCGCGAGACGAAAGGGCCTCGTGATACGCCTATTTTTATAGG
TTAATGTCATGATAATAATGGTTTCTTAGACGGATCGCTTGCCTGTAACTTACACGCGCCTCGTATCT
TTTAATGATGGAATAATTTGGGAATTTACTCTGTGTTTATTTATTTTTATGTTTTGTATTTGGATTTTA
GAAAGTAAATAAAGAAGGTAGAAGAGTTACGGAATGAAGAAAAAAAAATAAACAAAGGTTTAAAA
AATTTCAACAAAAAGCGTACTTTACATATATATTTATTAGACAAGAAAAGCAGATTAAATAGATATA
CATTCGATTAACGATAAGTAAAATGTAAAATCACAGGATTTTCGTGTGTGGTCTTCTACACAGACAA
GATGAAACAATTCGGCATTAATACCTGAGAGCAGGAAGAGCAAGATAAAAGGTAGTATTTGTTGGC
GATCCCCCTAGAGTCTTTTACATCTTCGGAAAACAAAAACTATTTTTTCTTTAATTTCTTTTTTTACTT
TCTATTTTTAATTTATATATTTATATTAAAAAATTTAAATTA

pND004

TGGGAAGCCCGATGCGCCAGAGTTGTTTCTGAAACATGGCAAAGGTAGCGTTGCCAATGATGTTAC
AGATGAGATGGTCAGACTAAACTGGCTGACGGAATTTATGCCTCTTCCGACCATCAAGCATTTTATC
CGTACTCCTGATGATGCATGGTTACTCACCACTGCGATCCCCGGGAAAACAGCATTCCAGGTATTAG
AAGAATATCCTGATTCAGGTGAAAATATTGTTGATGCGCTGGCAGTGTTCCTGCGCCGGTTGCATTC
GATTCCTGTTTGTAATTGTCCTTTTAACAGCGATCGCGTATTTCGTCTCGCTCAGGCGCAATCACGAA
TGAATAACGGTTTGGTTGATGCGAGTGATTTTGATGACGAGCGTAATGGCTGGCCTGTTGAACAAGT
CTGGAAAGAAATGCATAAACTTTTGCCATTCTCACCGGATTCAGTCGTCACTCATGGTGATTTCTCA
CTTGATAACCTTATTTTTGACGAGGGGAAATTAATAGGTTGTATTGATGTTGGACGAGTCGGAATCG
CAGACCGATACCAGGATCTTGCCATCCTATGGAACTGCCTCGGTGAGTTTTCTCCTTCATTACAGAA
ACGGCTTTTTCAAAAATATGGTATTGATAATCCTGATATGAATAAATTGCAGTTTCATTTGATGCTCG
ATGAGTTTTTCTAAGAATTAATTCATGAGCGGATACATATTTGAATGTATTTAGAAAAATAAACAAA
TAGGGGTTCCGCGCACATTTCCCCGAAAAGTGCCACCTAAATTGTAAGCGTTAATATTTTGTTAAAA
TTCGCGTTAAATTTTTGTTAAATCAGCTCATTTTTTAACCAATAGGCCGAAATCGGCAAAATCCCTTA
TAAATCAAAAGAATAGACCGAGATAGGGTTGAGTGTTGTTCCAGTTTGGAACAAGAGTCCACTATT
AAAGAACGTGGACTCCAACGTCAAAGGGCGAAAAACCGTCTATCAGGGCGATGGCCCACTACGTGA
ACCATCACCCTAATCAAGTTTTTTGGGGTCGAGGTGCCGTAAAGCACTAAATCGGAACCCTAAAGG
GAGCCCCCGATTTAGAGCTTGACGGGGAAAGCCGGCGAACGTGGCGAGAAAGGAAGGGAAGAAAG
CGAAAGGAGCGGGCGCTAGGGCGCTGGCAAGTGTAGCGGTCACGCTGCGCGTAACCACCACACCCG
CCGCGCTTACTACAGGGCGCGTCCCATTCGCTGAGGGCCAATCCGGATATAGTTCCAGCAAAAAAC
CCCTCAAGACCCGTTTAGAGGCCCCAAGGGGTTATGCTAGTTATTGCTCAGCGGTGGCAGCAGCCAA
CTCAGCTTCCTTGAGACCTATAAACGCAGAAAGGCCCACCCGAAGGTGAGCCAGTGTGACTCTAGT
AGAGAGCGTTCACCGACAAACAACAGATAAAACGAAAGGCCCAGTCTTTCGACTGAGCCTTTCGTT
TTATTTGATGCCTGGTCATTTGTACAGTTCATCCATACCATGCGTGATGCCCGCTGCGGTTACGAACT
CCAGCAGAACCATATGATCGCGTTTCTCGTTCGGATCTTTAGACAGAACGCTTTGCGTGCTCAGATA
GTGATTGTCTGGCAGCAGAACAGGACCATCACCGATTGGAGTGTTTTGCTGGTAGTGATCAGCCAGC
TGCACGCTGCCATCCTCCACGTTGTGGCGAATTTTAAAATTCGCTTTAATGCCATTTTTTTGTTTATC
GGCGGTGATGTAAACATTGTGGCTGTTAAAATTGTATTCCAGCTTATGGCCCAGGATATTGCCGTCC
TCTTTAAAGTCAATGCCTTTCAGCTCAATGCGGTTTACCAGGGTATCGCCTTCAAATTTCACTTCCGC
ACGCGTTTTGTACGTGCCGTCATCCTTAAAGGAAATCGTGCGTTCCTGCACATAGCCTTCCGGCATG
GCGGACTTGAAGAAGTCATGCTGCTTCATATGGTCCGGATAACGAGCAAAGCACTGAACACCATAA
GTCAGCGTCGTTACCAGAGTCGGCCAAGGAACCGGCAGTTTACCAGTAGTACAGATGAACTTCAGC
GTCAGTTTACCATTAGTTGCGTCACCTTCACCCTCGCCACGCACGGAAAACTTATGACCGTTGACAT
CACCATCCAGTTCCACCAGAATAGGGACGACACCAGTGAACAGCTCTTCGCCTTTACGCATCTAGTA
TTTCTCCTCTTTCTCTAGTAGCTAAGATCCGAACCTGCCGTTTCTTGAGTTGCCGCGATGTTAAGAAA
ACATTCATAAATTAAATGTGAATTGCCGCACACATTATTAAATAAGATTTACAAAATGTTCAAAATG
ACGCATGAAATCACGTTTCACTTTCGGTCTCAACCGCTGGCGGCGTTGATCACCGCAGTACGCACGG
CATACCAGAAAGCGGACATCTGCGGGATGTTCGGCATGATTTCACCTTTCTGGGCGTTTTCCATAGT
GGCGGCAATACGCGGATCTTTCACCAACTCTTCCTCGTAAGACTTCAGCGCTACGGCACCCAGCGGT
TTGTCTTTATTAACCGCTTCCAGACCTTCATCAGTCAGCAGATAGTTTTCGAGGAACTCTTTTGCCAG

CTCTTTGTTCGGACTGGCGGCGTTAATACCTGCGCTCAGCACGCCAACGAACGGTTTGGATGGTTGA
CCCTTGAAGGTCGGCAGTACCGTTACACCATAATTCACTTTGCTGGTGTCGATGTTGGACCATGCCC
ACGGGCCGTTGATGGTCATCGCTGTTTCGCCTTTATTAAAGGCAGCTTCTGCGATGGAGTAATCGGT
GTCTGCATTCATGTGTTTGTTTTTAATCAGGTCAACCAGGAAGGTCAGACCCGCTTTCGCGCCAGCG
TTATCCACGCCCACGTCTTTAATGTCGTACTTGCCGTTTTCATACTTGAACGCATAACCCCCGTCAGC
AGCAATCAGCGGCCAGGTGAAGTACGGTTCTTGCAGGTTGAACATCAGCGCGCTCTTACCTTTCGCT
TTCAGTTCTTTATCCAGCGCCGGGATCTCTTCCCAGGTTTTTGGCGGGTTCGGCAGCAGATCTTTGTT
ATAAATCAGCGATAACGCTTCAACAGCGATCGGGTAAGCAATCAGCTTGCCGTTGTAACGTACGGC
ATCCCAGGTAAACGGATACAGCTTGTCCTGGAACGCTTTGTCCGGGGTGATTTCAGCCAACAGGCCA
GATTGAGCGTAGCCACCAAAGCGGTCGTGTGCCCAGAAGATAATGTCAGGGCCATCGCCAGTTGCC
GCAACCTGTGGGAATTTCTCTTCCAGTTTATCCGGATGCTCAACGGTGACTTTAATTCCGGTATCTTT
CTCGAATTTCTTACCGACTTCAGCGAGTCCGTTATAGCCTTTATCGCCGTTAATCCAGATTACCAGTT
TACCTTCTTCGATTTTCATGCCGCTGCTGTGATGATGATGATGATGGCTGCTATTCATGGTATATCTC
CTTCTTAAAGTTAAACAAAATTATTTCTAGAGGGGAATTGTTATCCGCTCACAATTCCCCTATAGTG
AGTCGTATTAATTTCGCGGGATCGAGATCGATCTCGATCCTCTACGCCGGACGCATCGTGGCCGGCA
TCACCGGCGCCACAGGTGCGGTTGCTGGCGCCTATATCGCCGACATCACCGATGGGGAAGATCGGG
CTCGCCACTTCGGGCTCATGAGCGCTTGTTTCGGCGTGGGTATGGTGGCAGGCCCCGTGGCCGGGGG
ACTGTTGGGCGCCATCTCCTTGCATGCACCATTCCTTGCGGCGGCGGTGCTCAACGGCCTCAACCTA
CTACTGGGCTGCTTCCTAATGCAGGAGTCGCATAAGGGAGAGCGTCGAGATCCCGGACACCATCGA
ATGGCGCAAAACCTTTCGCGGTATGGCATGATAGCGCCCGGAAGAGAGTCAATTCAGGGTGGTGAA
TGTGAAACCAGTAACGTTATACGATGTCGCAGAGTATGCCGGTGTCTCTTATCAGACCGTTTCCCGC
GTGGTGAACCAGGCCAGCCACGTTTCTGCGAAAACGCGGGAAAAAGTGGAAGCGGCGATGGCGGA
GCTGAATTACATTCCCAACCGCGTGGCACAACAACTGGCGGGCAAACAGTCGTTGCTGATTGGCGTT
GCCACCTCCAGTCTGGCCCTGCACGCGCCGTCGCAAATTGTCGCGGCGATTAAATCTCGCGCCGATC
AACTGGGTGCCAGCGTGGTGGTGTCGATGGTAGAACGAAGCGGCGTCGAAGCCTGTAAAGCGGCGG
TGCACAATCTTCTCGCGCAACGCGTCAGTGGGCTGATCATTAACTATCCGCTGGATGACCAGGATGC
CATTGCTGTGGAAGCTGCCTGCACTAATGTTCCGGCGTTATTTCTTGATGTCTCTGACCAGACACCCA
TCAACAGTATTATTTTCTCCCATGAAGACGGTACGCGACTGGGCGTGGAGCATCTGGTCGCATTGGG
TCACCAGCAAATCGCGCTGTTAGCGGGCCCATTAAGTTCTGTCTCGGCGCGTCTGCGTCTGGCTGGC
TGGCATAAATATCTCACTCGCAATCAAATTCAGCCGATAGCGGAACGGGAAGGCGACTGGAGTGCC
ATGTCCGGTTTTCAACAAACCATGCAAATGCTGAATGAGGGCATCGTTCCCACTGCGATGCTGGTTG
CCAACGATCAGATGGCGCTGGGCGCAATGCGCGCCATTACCGAGTCCGGGCTGCGCGTTGGTGCGG
ACATCTCGGTAGTGGGATACGACGATACCGAAGACAGCTCATGTTATATCCCGCCGTTAACCACCAT
CAAACAGGATTTTCGCCTGCTGGGGCAAACCAGCGTGGACCGCTTGCTGCAACTCTCTCAGGGCCAG
GCGGTGAAGGGCAATCAGCTGTTGCCCGTCTCACTGGTGAAAAGAAAAACCACCCTGGCGCCCAAT
ACGCAAACCGCCTCTCCCCGCGCGTTGGCCGATTCATTAATGCAGCTGGCACGACAGGTTTCCCGAC
TGGAAAGCGGGCAGTGAGCGCAACGCAATTAATGTAAGTTAGCTCACTCATTAGGCACCGGGATCT
CGACCGATGCCCTTGAGAGCCTTCAACCCAGTCAGCTCCTTCCGGTGGGCGCGGGGCATGACTATCG
TCGCCGCACTTATGACTGTCTTCTTTATCATGCAACTCGTAGGACAGGTGCCGGCAGCGCTCTGGGT
CATTTTCGGCGAGGACCGCTTTCGCTGGAGCGCGACGATGATCGGCCTGTCGCTTGCGGTATTCGGA
ATCTTGCACGCCCTCGCTCAAGCCTTCGTCACTGGTCCCGCCACCAAACGTTTCGGCGAGAAGCAGG
CCATTATCGCCGGCATGGCGGCCCCACGGGTGCGCATGATCGTGCTCCTGTCGTTGAGGACCCGGCT
AGGCTGGCGGGGTTGCCTTACTGGTTAGCAGAATGAATCACCGATACGCGAGCGAACGTGAAGCGA
CTGCTGCTGCAAAACGTCTGCGACCTGAGCAACAACATGAATGGTCTTCGGTTTCCGTGTTTCGTAA
AGTCTGGAAACGCGGAAGTCAGCGCCCTGCACCATTATGTTCCGGATCTGCATCGCAGGATGCTGCT
GGCTACCCTGTGGAACACCTACATCTGTATTAACGAAGCGCTGGCATTGACCCTGAGTGATTTTTCT
CTGGTCCCGCCGCATCCATACCGCCAGTTGTTTACCCTCACAACGTTCCAGTAACCGGGCATGTTCA
TCATCAGTAACCCGTATCGTGAGCATCCTCTCTCGTTTCATCGGTATCATTACCCCCATGAACAGAA
ATCCCCCTTACACGGAGGCATCAGTGACCAAACAGGAAAAAACCGCCCTTAACATGGCCCGCTTTA
TCAGAAGCCAGACATTAACGCTTCTGGAGAAACTCAACGAGCTGGACGCGGATGAACAGGCAGACA
TCTGTGAATCGCTTCACGACCACGCTGATGAGCTTTACCGCAGCTGCCTCGCGCGTTTCGGTGATGA
CGGTGAAAACCTCTGACACATGCAGCTCCCGGAGACGGTCACAGCTTGTCTGTAAGCGGATGCCGG
GAGCAAACAAGCCCGTCAGGGCGCGTCAGCGGGTGTTGGCGGGTGTCGGGGCGCAGCCATGACCCA
GTCACGTAGCGATAGCGGAGTGTATACTGGCTTAACTATGCGGCATCAGAGCAGATTGTACTGAGA
GTGCACCATTGCGGTGTGAAATACCGCACAGATGCGTAAGGAGAAAATACCGCATCAGGCGCTCTT

CCGCTTCCTCGCTCACTGACTCGCTGCGCTCGGTCGTTCGGCTGCGGCGAGCGGTATCAGCTCACTC
AAAGGCGGTAATACGGTTATCCACAGAATCAGGGGATAACGCAGGAAAGAACATGTGAGCAAAAG
GCCAGCAAAAGGCCAGGAACCGTAAAAAGGCCGCGTTGCTGGCGTTTTTCCATAGGCTCCGCCCCC
CTGACGAGCATCACAAAAATCGACGCTCAAGTCAGAGGTGGCGAAACCCGACAGGACTATAAAGAT
ACCAGGCGTTTCCCCCTGGAAGCTCCCTCGTGCGCTCTCCTGTTCCGACCCTGCCGCTTACCGGATAC
CTGTCCGCCTTTCTCCCTTCGGGAAGCGTGGCGCTTTCTCATAGCTCACGCTGTAGGTATCTCAGTTC
GGTGTAGGTCGTTCGCTCCAAGCTGGGCTGTGTGCACGAACCCCCCGTTCAGCCCGACCGCTGCGCC
TTATCCGGTAACTATCGTCTTGAGTCCAACCCGGTAAGACACGACTTATCGCCACTGGCAGCAGCCA
CTGGTAACAGGATTAGCAGAGCGAGGTATGTAGGCGGTGCTACAGAGTTCTTGAAGTGGTGGCCTA
ACTACGGCTACACTAGAAGGACAGTATTTGGTATCTGCGCTCTGCTGAAGCCAGTTACCTTCGGAAA
AAGAGTTGGTAGCTCTTGATCCGGCAAACAAACCACCGCTGGTAGCGGTGGTTTTTTTGTTTGCAAG
CAGCAGATTACGCGCAGAAAAAAAGGATCTCAAGAAGATCCTTTGATCTTTTCTACGGGGTCTGAC
GCTCAGTGGAACGAAAACTCACGTTAAGGGATTTTGGTCATGAACAATAAAACTGTCTGCTTACATA
AACAGTAATACAAGGGGTGTTATGAGCCATATTCAACGGGAAACGTCTTGCTCTAGGCCGCGATTA
AATTCCAACATGGATGCTGATTTATATGGGTATAAATGGGCTCGCGATAATGTCGGGCAATCAGGTG
CGACAATCTATCGATTGTA

pND005

AGCAAACTGAAACGTTTTCATCGCTCTGGAGTGAATACCACGACGATTTCCGGCAGTTTCTACACAT
ATATTCGCAAGATGTGGCGTGTTACGGTGAAAACCTGGCCTATTTCCCTAAAGGGTTTATTGAGAAT
ATGTTTTTCGTCTCAGCCAATCCCTGGGTGAGTTTCACCAGTTTTGATTTAAACGTGGCCAATATGGA
CAACTTCTTCGCCCCCGTTTTCACCATGGGCAAATATTATACGCAAGGCGACAAGGTGCTGATGCCG
CTGGCGATTCAGGTTCATCATGCCGTTTGTGATGGCTTCCATGTCGGCAGAATGCTTAATGAATTAC
AACAGTACTGCGATGAGTGGCAGGGCGGGGCGTAATTTTTTTAAGGCAGTTATTGGTGCCCTTAAAC
GCCTGGTTGCTACGCCTGAATAAGTGATAATAAGCGGATGAATGGCAGAAATTCGAAAGCAAATTC
GACCCGGTCGTCGGTTCAGGGCAGGGTCGTTAAATAGCCGCTTATGTCTATTGCTGGTTTACCGGTT
TATTGACTACCGGAAGCAGTGTGACCGTGTGCTTCTCAAATGCCTGAGGCCAGTTTGCTCAGGCTCT
CCCCGTGGAGGCTAAGAAACCATTATTATCATGACATTAACCTATAAAAATAGGCGTATCACGAGG
CCCTTTCGTCTCGCGCGTTTCGGTGATGACGGTGAAAACCTCTGACACATGCAGCTCCCGGAGACGG
TCACAGCTTGTCTGTAAGCGGATGCCGGGAGCAGACAAGCCCGTCAGGGCGCGTCAGCGGGTGTTG
GCGGGTGTCGGGGCTGGCTTAACTATGCGGCATCAGAGCAGATTGTACTGAGAGTGCACCATAACG
CATTTAAGCATAAACACGCACTATGCCGTTCTTCTCATGTATATATATATACAGGCAACACGCAGAT
ATAGGTGCGACGTGAACAGTGAGCTGTATGTGCGCAGCTCGCGTTGCATTTTCGGAAGCGCTCGTTT
TCGGAAACGCTTTGAAGTTCCTATTCCGAAGTTCCTATTCTCTAGCTAGAAAGTATAGGAACTTCAG
AGCGCTTTTGAAAACCAAAAGCGCTCTGAAGACGCACTTTCAAAAAACCAAAAACGCACCGGACTG
TAACGAGCTACTAAAATATTGCGAATACCGCTTCCACAAACATTGCTCAAAAGTATCTCTTTGCTAT
ATATCTCTGTGCTATATCCCTATATAACCTACCCATCCACCTTTCGCTCCTTGAACTTGCATCTAAAC
TCGACCTCTACATTTTTTATGTTTATCTCTAGTATTACTCTTTAGACAAAAAAATTGTAGTAAGAACT
ATTCATAGAGTGAATCGAAAACAATACGAAAATGTAAACATTTCCTATACGTAGTATATAGAGACA
AAATAGAAGAAACCGTTCATAATTTTCTGACCAATGAAGAATCATCAACGCTATCACTTTCTGTTCA
CAAAGTATGCGCAATCCACATCGGTATAGAATATAATCGGGGATGCCTTTATCTTGAAAAAATGCAC
CCGCAGCTTCGCTAGTAATCAGTAAACGCGGGAAGTGGAGTCAGGCTTTTTTTATGGAAGAGAAAA
TAGACACCAAAGTAGCCTTCTTCTAACCTTAACGGACCTACAGTGCAAAAAGTTATCAAGAGACTGC
ATTATAGAGCGCACAAAGGAGAAAAAAAGTAATCTAAGATGCTTTGTTAGAAAAATAGCGCTCTCG
GGATGCATTTTTGTAGAACAAAAAAGAAGTATAGATTCTTTGTTGGTAAAATAGCGCTCTCGCGTTG
CATTTCTGTTCTGTAAAAATGCAGCTCAGATTCTTTGTTTGAAAAATTAGCGCTCTCGCGTTGCATTT
TTGTTTTACAAAAATGAAGCACAGATTCTTCGTTGGTAAAATAGCGCTTTCGCGTTGCATTTCTGTTC
TGTAAAAATGCAGCTCAGATTCTTTGTTTGAAAAATTAGCGCTCTCGCGTTGCATTTTTGTTCTACAA
AATGAAGCACAGATGCTTCGTTGCTTGCATGCAACTTCTTTTCTTTTTTTTTCTTTTCTCTCTCCCCCG
TTGTTGTCTCACCATATCCGCAATGACAAAAAAATGATGGAAGACACTAAAGGAAAAAATTAACGA
CAAAGACAGCACCAACAGATGTCGTTGTTCCAGAGCTGATGAGGGGTATCTCGAAGCACACGAAAC
TTTTTCCTTCCTTCATTCACGCACACTACTCTCTAATGAGCAACGGTATACGGCCTTCCTTCCAGTTA
CTTGAATTTGAAATAAAAAAAAGTTTGCTGTCTTGCTATCAAGTATAAATAGACCTGCAATTATTAA
TCTTTTGTTTCCTCGTCATTGTTCTCGTTCCCTTTCTTCCTTGTTTCTTTTTCTGCACAATATTTCAAGC
TATACCAAGCATACAATCAACTCCAAGCTTGAAGCAAGCCTCCTTGAGACCGAAAGTGAAACGTGA

TTTCATGCGTCATTTTGAACATTTTGTAAATCTTATTTAATAATGTGTGCGGCAATTCACATTTAATTT
ATGAATGTTTTCTTAACATCGCGGCAACTCAAGAAACGGCAGGTTCGGATCTTAGCTACTAGAGAAA
GAGGAGAAATACTAGATGCGTAAAGGCGAAGAGCTGTTCACTGGTGTCGTCCCTATTCTGGTGGAA
CTGGATGGTGATGTCAACGGTCATAAGTTTTCCGTGCGTGGCGAGGGTGAAGGTGACGCAACTAAT
GGTAAACTGACGCTGAAGTTCATCTGTACTACTGGTAAACTGCCGGTTCCTTGGCCGACTCTGGTAA
CGACGCTGACTTATGGTGTTCAGTGCTTTGCTCGTTATCCGGACCATATGAAGCAGCATGACTTCTTC
AAGTCCGCCATGCCGGAAGGCTATGTGCAGGAACGCACGATTTCCTTTAAGGATGACGGCACGTAC
AAAACGCGTGCGGAAGTGAAATTTGAAGGCGATACCCTGGTAAACCGCATTGAGCTGAAAGGCATT
GACTTTAAAGAGGACGGCAATATCCTGGGCCATAAGCTGGAATACAATTTTAACAGCCACAATGTTT
ACATCACCGCCGATAAACAAAAAAATGGCATTAAAGCGAATTTTAAAATTCGCCACAACGTGGAGG
ATGGCAGCGTGCAGCTGGCTGATCACTACCAGCAAAACACTCCAATCGGTGATGGTCCTGTTCTGCT
GCCAGACAATCACTATCTGAGCACGCAAAGCGTTCTGTCTAAAGATCCGAACGAGAAACGCGATCA
TATGGTTCTGCTGGAGTTCGTAACCGCAGCGGGCATCACGCATGGTATGGATGAACTGTACAAATGA
CCAGGCATCAAATAAAACGAAAGGCTCAGTCGAAAGACTGGGCCTTTCGTTTTATCTGTTGTTTGTC
GGTGAACGCTCTCTACTAGAGTCACACTGGCTCACCTTCGGGTGGGCCTTTCTGCGTTTATAGGTCTC
AATTCCGGGCGAATTTCTTATGATTTATGATTTTTATTATTAAATAAGTTATAAAAAAAATAAGTGTA
TACAAATTTTAAAGTGACTCTTAGGTTTTAAAACGAAAATTCTTATTCTTGAGTAACTCTTTCCTGTA
GGTCAGGTTGCTTTCTCAGGTATAGCATGAGGTCGCTCTTATTGACCACACCTCTACCGGCATGCCG
GCAAGTGCACAAACAATACTTAAATAAATACTACTCAGTAATAACCTATTTCTTAGCATTTTTGACG
AAATTTGCTATTTTGTTAGAGTCTTTTACACCATTTGTCTCCACACCTCCGCTTACATCAACACCAAT
AACGCCATTTAATCTAAGCGCATCACCAACATTTTCTGGCGTCAGTCCACCAGCTAACATAAAATGT
AAGCTTTCGGGGCTCTCTTGCCTTCCAACCCAGTCAGAAATCGAGTTCCAATCCAAAAGTTCACCTG
TCCCACCTGCTTCTGAATCAAACAAGGGAATAAACGAATGAGGTTTCTGTGAAGCTGCACTGAGTA
GTATGTTGCAGTCTTTTGGAAATACGAGTCTTTTAATAACTGGCAAACCGAGGAACTCTTGGTATTC
TTGCCACGACTCATCTCCATGCAGTTGGACGATATCAATGCCGTAATCATTGACCAGAGCCAAAACA
TCCTCCTTAGGTTGATTACGAAACACGCCAACCAAGTATTTCGGAGTGCCTGAACTATTTTTATATGC
TTTTACAAGACTTGAAATTTTCCTTGCAATAACCGGGTCAATTGTTCTCTTTCTATTGGGCACACATA
TAATACCCAGCAAGTCAGCATCGGAATCTAGAGCACATTCTGCGGCCTCTGTGCTCTGCAAGCCGCA
AACTTTCACCAATGGACCAGAACTACCTGTGAAATTAATAACAGACATACTCCAAGCTGCCTTTGTG
TGCTTAATCACGTATACTCACGTGCTCAATAGTCACCAATGCCCTCCCTCTTGGCCCTCTCCTTTTCTT
TTTTCGACCGAATTAATTCGTAATCATGTCATAGCTGTTTCCTGTGTGAAATTGTTATCCGCTCACAA
TTCCACACAACATACGAGCCGGAAGCATAAAGTGTAAAGCCTGGGGTGCCTAATGAGTGAGCTAAC
TCACATTAATTGCGTTGCGCTCACTGCCCGCTTTCCAGTCGGGAAACCTGTCGTGCCAGCTGGTTCTT
TCCGCCTCAGGACTCTTCCTTTTTCAATATTATTGAAGCATTTATCAGGGTTATTGTCTCATGAGCGG
ATACATATTTGAATGTATTTAGAAAAATAAACAAATAGGGGTTCCGCGCACATTTCCCCGAAAAGTG
CCACCTGACGCGCCCTGTAGCGGCGCATTAAGCGCGGCGGGTGTGGTGGTTACGCGCAGCGTGACC
GCTACACTTGCCAGCGCCCTAGCGCCCGCTCCTTTCGCTTTCTTCCCTTCCTTTCTCGCCACGTTCGCC
GGCTTTCCCCGTCAAGCTCTAAATCGGGGGCTCCCTTTAGGGTTCCGATTTAGTGCTTTACGGCACCT
CGACCCCAAAAAACTTGATTAGGGTGATGGTTCACGTAGTGGGCCATCGCCCTGATAGACGGTTTTT
CGCCCTTTGACGTTGGAGTCCACGTTCTTTAATAGTGGACTCTTGTTCCAAACTGGAACAACACTCA
ACCCTATCTCGGTCTATTCTTTTGATTTATAAGGGATTTTGCCGATTTCGGCCTATTGGTTAAAAAAT
GAGCTGATTTAACAAAAATTTAACGCGAATTTTAACAAAATATTAACGCTTACAATTTACGCGCCTA
AATTATGCGGCCGCATATAACTGCATTAATGAATCGGCCAACGCGCGGGGAGAGGCGGTTTGCGTA
TTGGGCGCTCTTCCGCTTCCTCGCTCACTGACTCGCTGCGCTCGGTCGTTCGGCTGCGGCGAGCGGTA
TCAGCTCACTCAAAGGCGGTAATACGGTTATCCACAGAATCAGGGGATAACGCAGGAAAGAACATG
TGAGCAAAAGGCCAGCAAAAGGCCAGGAACCGTAAAAAGGCCGCGTTGCTGGCGTTTTTCCATAGG
CTCCGCCCCCCTGACGAGCATCACAAAAATCGACGCTCAAGTCAGAGGTGGCGAAACCCGACAGGA
CTATAAAGATACCAGGCGTTTCCCCCTGGAAGCTCCCTCGTGCGCTCTCCTGTTCCGACCCTGCCGCT
TACCGGATACCTGTCCGCCTTTCTCCCTTCGGGAAGCGTGGCGCTTTCTCATAGCTCACGCTGTAGGT
ATCTCAGTTCGGTGTAGGTCGTTCGCTCCAAGCTGGGCTGTGTGCACGAACCCCCCGTTCAGCCCGA
CCGCTGCGCCTTATCCGGTAACTATCGTCTTGAGTCCAACCCGGTAAGACACGACTTATCGCCACTG
GCAGCAGCCACTGGTAACAGGATTAGCAGAGCGAGGTATGTAGGCGGTGCTACAGAGTTCTTGAAG
TGGTGGCCTAACTACGGCTACACTAGAAGAACAGTATTTGGTATCTGCGCTCTGCTGAAGCCAGTTA
CCTTCGGAAAAAGAGTTGGTAGCTCTTGATCCGGCAAACAAACCACCGCTGGTAGCGGTGGTTTTTT
TGTTTGCAAGCAGCAGATTACGCGCAGAAAAAAAGGATCTCAAGAAGATCCTTTGATCTTTTCTACG

GGGTCTGACGCTCAGTGGAACGAAAACTCACGTTAAGGGATTTTGGTCATGAGATTATCAAAAAGG
ATCTTCACCTAGATCCTTTTAAATTAAAAATGAAGTTTTAAATCAATCTAAAGTATATATGAGTAAA
CTTGGTCTGACAGTTACCAATGCTTAATCAGTGAGGCACCTATCTCAGCGATCTGTCTATTTCGTTCA
TCCATAGTTGCCTGACTCCCCGTCGTGTAGATAACTACGATACGGGAGGGCTTACCATCTGGCCCCA
GTGCTGCAATGATACCGCGAGATCCACGCTCACCGGCTCCAGATTTATCAGCAATAAACCAGCCAG
CCGCCTGTGACGGAAGATCACTTCGCAGAATAAATAAATCCTGGTGTCCCTGTTGATACCGGGAAGC
CCTGGGCCAACTTTTGGCGAAAATGAGACGTTGATCGGCACGTAAGAGGTTCCAACTTTCACCATAA
TGAAATAAGATCACTACCGGGCGTATTTTTTGAGTTGTCGAGATTTTCAGGAGCTAAGGAAGCTAAA
ATGGAGAAAAAAATCACTGGATATACCACCGTTGATATATCCCAATGGCATCGTAAAGAACATTTT
GAGGCATTTCAGTCAGTTGCTCAATGTACCTATAACCAGACCGTTCAGCTGGATATTACGGCCTTTTT
AAAGACCGTAAAGAAAAATAAGCACAAGTTTTATCCGGCCTTTATTCACATTCTTGCCCGCCTGATG
AATGCTCATCCGGAATTACGTATGGCAATGAAAGACGGTGAGCTGGTGATATGGGATAGTGTTCAC
CCTTGTTACACCGTTTTCCATG

pIFU037

ATTGTGAGCGGATAACAATTTCACACAGGAAACAGCTATGACCATGATTACGCCAAGCTCGAAATT
AACCCTCACTAAAGGGAACAAAAGCTGGTACCAATTCCTTGAATTTTCAAAAATTCTTACTTTTTTT
TGGATGGACGCAAAGAAGTTTAATAATCATATTACATGGCATTACCACCATATACATATCCATATCT
AATCTTACTTATATGTTGTGGAAATGTAAAGAGCCCCATTATCTTAGCCTAAAAAAACCTTCTCTTTG
GAACTTTCAGTAATACGCTTAACTGCTCATTGCTATATTGAAGTACGGATTAGAAGCCGCCGAGCGG
GTGACAGCCCTCCGAAGGAAGACTCTCCTCCGTGCGTCCTCGTCTTCACCGGTCGCGTTCCTGAAAC
GCAGATGTGCCTCGCGCCGCACTGCTCCGAACAATAAAGATTCTACAATACTAGCTTTTATGGTTAT
GAAGAGGAAAAATTGGCAGTAACCTGGCCCCACAAACCTTCAAATGAACGAATCAAATTAACAACC
ATAGGATGATAATGCGATTAGTTTTTTAGCCTTATTTCTGGGGTAATTAATCAGCGAAGCGATGATT
TTTGATCTATTAACAGATATATAAATGCAAAAACTGCATAACCACTTTAACTAATACTTTCAACATTT
TCGGTTTGTATTACTTCTTATTCAAATGTAATAAAAGTATCAACAAAAAATTGTTAATATACCTCTAT
ACTTTAACGTCAAGGAGAAAAAACCCCGGATCGAATTCCCTACTTCATACATTTTCAATTAAGATGC
AGTTACTTCGCTGTTTTTCAATATTTTCTGTTATTGCTTCAGTTTTAGCACAGGAACTGACAACTATA
TGCGAGCAAATCCCCTCACCAACTTTAGAATCGACGCCGTACTCTTTGTCAACGACTACTATTTTGG
CCAACGGGAAGGCAATGCAAGGAGTTTTTGAATATTACAAATCAGTAACGTTTGTCAGTAATTGCG
GTTCTCACCCCTCAACAACTAGCAAAGGCAGCCCCATAAACACACAGTATGTTTTTAAGGACAATAG
CTCGACGATTGAAGGTAGATACCCATACGACGTTCCAGACTACGCTCTGCAGGCTAGTGGTGGAGG
AGGCTCTGGTGGAGGCGGTAGCGGAGGCGGAGGGTCGGCTAGCCATACAAACTTGTGCCCTTTTGG
TGAAGTTTTTAACGCCACCAGATTTGCATCTGTTTATGCTTGGAACAGGAAGAGAATCAGCAACTGT
GTTGCTGATTATTCTGTCCTATATAATTCCGCATCATTTTCCACTTTTAAGTGTTATGGAGTGTCTCCT
ACTAAATTAAATGATCTCTGCTTTACTAATGTCTATGCAGATTCATTTGAGAGACCACGGCCACAGC
TAACACCACGTCGTCCCTATCTGCTGCCCTAGGTCTATGAGTGGTTGCTGGATAACTTTACGGGCAT
GCATAAGGCTCGTAATATATATTCAGGGAGGCCACAACGGTTTCCCTCTACAAATAATTTTGTTTAA
CTTTCTAGAAATAATTTTGTTTAACTTTAAGAAGGAGATATACATATGCGTAAAGGAGAAGAACTTT
TCACTGGAGTTGTCCCAATTCTTGTTGAATTAGATGGTGATGTTAATGGGCACAAATTTTCTGTCAGT
GGAGAGGGTGAAGGTGATGCAACATACGGAAAACTTACCCTTAAATTTATTTGCACTACTGGAAAA
CTACCTGTTCCATGGCCAACACTTGTCACTACTTTCGGTTATGGTGTTCAATGCTTTGCGAGATACCC
AGATCATATGAAACAGCATGACTTTTTCAAGAGTGCCATGCCCGAAGGTTATGTACAGGAAAGAAC
TATATTTTTCAAAGATGACGGGAACTACAAGACACGTGCTGAAGTCAAGTTTGAAGGTGATACCCTT
GTTAATAGAATCGAGTTAAAAGGTATTGATTTTAAAGAAGATGGAAACATTCTTGGACACAAATTG
GAATACAACTATAACTCACACAATGTATACATCATGGCAGACAAACAAAAGAATGGAATCAAAGTT
AACTTCAAAATTAGACACAACATTGAAGATGGAAGCGTTCAACTAGCAGACCATTATCAACAAAAT
ACTCCAATTGGCGATGGCCCTGTCCTTTTACCAGACAACCATTACCTGTCCACACAATCTGCCCTTTC
GAAAGATCCCAACGAAAAGAGGGACCACATGGTCCTTCTTGAGTTTGTAACAGCTGCTGGGATTAC
ACATGGCATGGATGAACTATACAAACTAGAGTGAGATCCGGCTGCTAACAAAGCCCGAAAGGAAGC
TGAGTTGGCTGCTGCCACCGCTGAGCAATAACTAGCATAACCCCTTGGGGCCTCTAAACGGGTCTTG
AGGGGTTTTTTGCTGAAAGGAGGAACTATATCCGGATGGTCTCAAGAGTAGTAGTACTTTCTTTTGA
ACTTCTACATGCACCAGCAACTGTTTGTGGACCTAAAAAGTCTACTAATTTGGTTAAAAACAAACTC
GAGGGGGGCGGATCCGAACAAAAGCTTATTTCTGAAGAGGACTTGTAATAGAGATCTGATAACAAC
AGTGTAGATGTAACAAAATCGACTTTGTTCCCACTGTACTTTTAGCTCGTACAAAATACAATATACT

TTTCATTTCTCCGTAAACAACATGTTTTCCCATGTAATATCCTTTTCTATTTTTCGTTCCGTTACCAAC
TTTACACATACTTTATATAGCTATTCACTTCTATACACTAAAAAACTAAGACAATTTTAATTTTGCTG
CCTGCCATATTTCAATTTGTTATAAATTCCTATAATTTATCCTATTAGTAGCTAAAAAAAGATGAATG
TGAATCGAATCCTAAGAGAATTGAGCTCCAATTCGCCCTATAGTGAGTCGTATTACAATTCACTGGC
CGTCGTTTTACAACGTCGTGACTGGGAAAACCCTGGCGTTACCCAACTTAATCGCCTTGCAGCACAT
CCCCCTTTCGCCAGCTGGCGTAATAGCGAAGAGGCCCGCACCGATCGCCTTTCCCAACAGTTGCGCA
GCCTGAATGGCGAATGGACGCGCCCTGTAGCGGCGCATTAAGCGCGGCGGGTGTGGTGGTTACGCG
CAGCGTGACCGCTACACTTGCCAGCGCCCTAGCGCCCGCTCCTTTCGCTTTCTTCCCTTCCTTTCTCG
CCACGTTCGCCGGCTTTCCCCGTCAAGCTCTAAATCGGGGGCTCCCTTTAGGGTTCCGATTTAGTGCT
TTACGGCACCTCGACCCCAAAAAACTTGATTAGGGTGATGGTTCACGTAGTGGGCCATCGCCCTGAT
AGACGGTTTTTCGCCCTTTGACGTTGGAGTCCACGTTCTTTAATAGTGGACTCTTGTTCCAAACTGGA
ACAACACTCAACCCTATCTCGGTCTATTCTTTTGATTTATAAGGGATTTTGCCGATTTCGGCCTATTG
GTTAAAAAATGAGCTGATTTAACAAAAATTTAACGCGAATTTTAACAAAATATTAACGCTTACAATT
TCCTGATGCGGTATTTTCTCCTTACGCATCTGTGCGGTATTTCACACCGCATAGATCGGCAAGTGCAC
AAACAATACTTAAATAAATACTACTCAGTAATAACCTATTTCTTAGCATTTTTGACGAAATTTGCTAT
TTTGTTAGAGTCTTTTACACCATTTGTCTCCACACCTCCGCTTACATCAACACCAATAACGCCATTTA
ATCTAAGCGCATCACCAACATTTTCTGGCGTCAGTCCACCAGCTAACATAAAATGTAAGCTTTCGGG
GCTCTCTTGCCTTCCAACCCAGTCAGAAATCGAGTTCCAATCCAAAAGTTCACCTGTCCCACCTGCTT
CTGAATCAAACAAGGGAATAAACGAATGAGGTTTCTGTGAAGCTGCACTGAGTAGTATGTTGCAGT
CTTTTGGAAATACGAGTCTTTTAATAACTGGCAAACCGAGGAACTCTTGGTATTCTTGCCACGACTC
ATCTCCATGCAGTTGGACGATATCAATGCCGTAATCATTGACCAGAGCCAAAACATCCTCCTTAGGT
TGATTACGAAACACGCCAACCAAGTATTTCGGAGTGCCTGAACTATTTTTATATGCTTTTACAAGAC
TTGAAATTTTCCTTGCAATAACCGGGTCAATTGTTCTCTTTCTATTGGGCACACATATAATACCCAGC
AAGTCAGCATCGGAATCTAGAGCACATTCTGCGGCCTCTGTGCTCTGCAAGCCGCAAACTTTCACCA
ATGGACCAGAACTACCTGTGAAATTAATAACAGACATACTCCAAGCTGCCTTTGTGTGCTTAATCAC
GTATACTCACGTGCTCAATAGTCACCAATGCCCTCCCTCTTGGCCCTCTCCTTTTCTTTTTTCGACCGA
ATTAATTCTTAATCGGCAAAAAAAGAAAAGCTCCGGATCAAGATTGTACGTAAGGTGACAAGCTAT
TTTTCAATAAAGAATATCTTCCACTACTGCCATCTGGCGTCATAACTGCAAAGTACACATATATTAC
GATGCTGTTCTATTAAATGCTTCCTATATTATATATATATAGTAATGTCGTGATCTATGGTGCACTCTCA
GTACAATCTGCTCTGATGCCGCATAGTTAAGCCAGCCCCGACACCCGCCAACACCCGCTGACGCGCC
CTGACGGGCTTGTCTGCTCCCGGCATCCGCTTACAGACAAGCTGTGACCGTCTCCGGGAGCTGCATG
TGTCAGAGGTTTTCACCGTCATCACCGAAACGCGCGAGACGAAAGGGCCTCGTGATACGCCTATTTT
TATAGGTTAATGTCATGATAATAATGGTTTCTTAGACGGATCGCTTGCCTGTAACTTACACGCGCCTC
GTATCTTTTAATGATGGAATAATTTGGGAATTTACTCTGTGTTTATTTATTTTTATGTTTTGTATTTGG
ATTTTAGAAAGTAAATAAAGAAGGTAGAAGAGTTACGGAATGAAGAAAAAAAAAATAAACAAAGGT
TTAAAAAATTTCAACAAAAAGCGTACTTTACATATATATTTATTAGACAAGAAAAGCAGATTAAATA
GATATACATTCGATTAACGATAAGTAAAATGTAAAATCACAGGATTTTCGTGTGTGGTCTTCTACAC
AGACAAGATGAAACAATTCGGCATTAATACCTGAGAGCAGGAAGAGCAAGATAAAAGGTAGTATTT
GTTGGCGATCCCCCTAGAGTCTTTTACATCTTCGGAAAACAAAAACTATTTTTTCTTTAATTTCTTTTT
TTACTTTCTATTTTTAATTTATATATTTATATTAAAAAATTTAAATTATAATTATTTTTATAGCACGTG
ATGAAAAGGACCCAGGTGGCACTTTTCGGGGAAATGTGCGCGGAACCCCTATTTGTTTATTTTTCTA
AATACATTCAAATATGTATCCGCTCATGAGACAATAACCCTGATAAATGCTTCAATAATATTGAAAA
AGGAAGAGTATGAGCCATATTCAACGGGAAACGTCTTGCTCTAGGCCGCGATTAAATTCCAACATG
GATGCTGATTTATATGGGTATAAATGGGCTCGCGATAATGTCGGGCAATCAGGTGCGACAATCTATC
GATTGTATGGGAAGCCCGATGCGCCAGAGTTGTTTCTGAAACATGGCAAAGGTAGCGTTGCCAATG
ATGTTACAGATGAGATGGTCAGACTAAACTGGCTGACGGAATTTATGCCTCTTCCGACCATCAAGCA
TTTTATCCGTACTCCTGATGATGCATGGTTACTCACCACTGCGATCCCCGGGAAAACAGCATTCCAG
GTATTAGAAGAATATCCTGATTCAGGTGAAAATATTGTTGATGCGCTGGCAGTGTTCCTGCGCCGGT
TGCATTCGATTCCTGTTTGTAATTGTCCTTTTAACAGCGATCGCGTATTTCGTCTCGCTCAGGCGCAA
TCACGAATGAATAACGGTTTGGTTGATGCGAGTGATTTTGATGACGAGCGTAATGGCTGGCCTGTTG
AACAAGTCTGGAAAGAAATGCATAAACTTTTGCCATTCTCACCGGATTCAGTCGTCACTCATGGTGA
TTTCTCACTTGATAACCTTATTTTTGACGAGGGGAAATTAATAGGTTGTATTGATGTTGGACGAGTCG
GAATCGCAGACCGATACCAGGATCTTGCCATCCTATGGAACTGCCTCGGTGAGTTTTCTCCTTCATT
ACAGAAACGGCTTTTTCAAAAATATGGTATTGATAATCCTGATATGAATAAATTGCAGTTTCATTTG
ATGCTCGATGAGTTTTTCTAACCTCAGCAACTGTCAGACCAAGTTTACTCATATATACTTTAGATTGA

TTTAAAACTTCATTTTTAATTTAAAAGGATCTAGGTGAAGATCCTTTTTGATAATCTCATGACCAAAA
TCCCTTAACGTGAGTTTTCGTTCCACTGAGCGTCAGACCCCGTAGAAAAGATCAAAGGATCTTCTTG
AGATCCTTTTTTTCTGCGCGTAATCTGCTGCTTGCAAACAAAAAAACCACCGCTACCAGCGGTGGTT
TGTTTGCCGGATCAAGAGCTACCAACTCTTTTTCCGAAGGTAACTGGCTTCAGCAGAGCGCAGATAC
CAAATACTGTTCTTCTAGTGTAGCCGTAGTTAGGCCACCACTTCAAGAACTCTGTAGCACCGCCTAC
ATACCTCGCTCTGCTAATCCTGTTACCAGTGGCTGCTGCCAGTGGCGATAAGTCGTGTCTTACCGGG
TTGGACTCAAGACGATAGTTACCGGATAAGGCGCAGCGGTCGGGCTGAACGGGGGGTTCGTGCACA
CAGCCCAGCTTGGAGCGAACGACCTACACCGAACTGAGATACCTACAGCGTGAGCTATGAGAAAGC
GCCACGCTTCCCGAAGGGAGAAAGGCGGACAGGTATCCGGTAAGCGGCAGGGTCGGAACAGGAGA
GCGCACGAGGGAGCTTCCAGGGGGAAACGCCTGGTATCTTTATAGTCCTGTCGGGTTTCGCCACCTC
TGACTTGAGCGTCGATTTTTGTGATGCTCGTCAGGGGGGCGGAGCCTATGGAAAAACGCCAGCAAC
GCGGCCTTTTTACGGTTCCTGGCCTTTTGCTGGCCTTTTGCTCACATGTTCTTTCCTGCGTTATCCCCT
GATTCTGTGGATAACCGTATTACCGCCTTTGAGTGAGCTGATACCGCTCGCCGCAGCCGAACGACCG
AGCGCAGCGAGTCAGTGAGCGAGGAAGCGGAAGAGCGCCCAATACGCAAACCGCCTCTCCCCGCG
CGTTGGCCGATTCATTAATGCAGCTGGCACGACAGGTTTCCCGACTGGAAAGCGGGCAGTGAGCGC
AACGCAATTAATGTGAGTTAGCTCACTCATTAGGCACCCCAGGCTTTACACTTTATGCTTCCGGCTC
GTATGTTGTGTGGA

pZB558

ATTTCCTGGAAGATGCCAGGAAGATACTTAACAGGGAAGTGAGAGGGCCGCGGCAAAGCCGTTTTT
CCATAGGCTCCGCCCCCCTGACAAGCATCACGAAATCTGACGCTCAAATCAGTGGTGGCGAAACCC
GACAGGACTATAAAGATACCAGGCGTTTCCCCCTGGCGGCTCCCTCGTGCGCTCTCCTGTTCCTGCC
TTTCGGTTTACCGGTGTCATTCCGCTGTTATGGCCGCGTTTGTCTCATTCCACGCCTGACACTCAGTT
CCGGGTAGGCAGTTCGCTCCAAGCTGGACTGTATGCACGAACCCCCCGTTCAGTCCGACCGCTGCGC
CTTATCCGGTAACTATCGTCTTGAGTCCAACCCGGAAAGACATGCAAAAGCACCACTGGCAGCAGC
CACTGGTAATTGATTTAGAGGAGTTAGTCTTGAAGTCATGCGCCGGTTAAGGCTAAACTGAAAGGA
CAAGTTTTGGTGACTGCGCTCCTCCAAGCCAGTTACCTCGGTTCAAAGAGTTGGTAGCTCAGAGAAC
CTTCGAAAAACCGCCCTGCAAGGCGGTTTTTTCGTTTTCAGAGCAAGAGATTACGCGCAGACCAAAA
CGATCTCAAGAAGATCATCTTATTAATCAGATAAAATATTTCTAGATTTCAGTGCAATTTATCTCTTC
AAATGTAGCACCTGAAGTCAGCCCCATACGATATAAGTTGTAATTCTCATGTTTGACAGCTTATCAT
CGATAAGCTGGACGTCTAAGAAACCATTATTATCATGACATTAACCTATAAAAATAGGCGTATCACG
AGGCCCTTTGGATAACCAGAAGCAATAAAAAATCAAATCGGATTTCACTATATAATCTCACTTTATC
TAAGATGAATCCGATGGAAGCATCCTGTTTTCTCTCAATTTTTTTATCTAAAACCCAGCGTTCGATGC
TTCTTTGAGCGAACGATCAAAAATAAGTGCCTTCCCATCAAAAAAATATTGACAACATAAAAAACTT
TGTGTTATACTTGTAACGCTACATGGAGATTAACTCAATCTAGCTAGAGAGGCTTTACACTTTATGCT
TCCGGCTCGTATAATGTGTGGAATTGTGAGCGGATAACAATTTCACACAGGAAACAGCTATGACCAT
GATTACGGATTCACTGGAACTCTAGACCAAAGAGAGGACACAATGAACACGATTAACATCGCTAAG
AACGACTTCTCTGACATCGAACTGGCTGCTATCCCGTTCAACACTCTGGCTGACCATTACGGTGAGC
GTTTAGCTCGCGAACAGTTGGCCCTTGAGCATGAGTCTTACGAGATGGGTGAAGCACGCTTCCGCAA
GATGTTTGAGCGTCAACTTAAAGCTGGTGAGGTTGCGGATAACGCTGCCGCCAAGCCTCTCATCACT
ACCCTACTCCCTAAGATGATTGCACGCATCAACGACTGGTTTGAGGAAGTGAAAGCTAAGCGCGGC
AAGCGCCCGACAGCCTTCCAGTTCCTGCAAGAAATCAAGCCGGAAGCCGTAGCGTACATCACCATT
AAGACCACTCTGGCTTGCCTAACCAGTGCTGACAATACAACCGTTCAGGCTGTAGCAAGCGCAATC
GGTCGGGCCATTGAGGACGAGGCTCGCTTCGGTCGTATCCGTGACCTTGAAGCTAAGCACTTCAAGA
AAAACGTTGAGGAACAACTCAACAAGCGCGTAGGGCACGTCTACAAGAAAGCATTTATGCAAGTTG
TCGAGGCTGACATGCTCTCTAAGGGTCTACTCGGTGGCGAGGCGTGGTCTTCGTGGCATAAGGAAG
ACTCTATTCATGTAGGAGTACGCTGCATCGAGATGCTCATTGAGTCAACCGGAATGGTTAGCTTACA
CCGCCAAAATGCTGGCGTAGTAGGTCAAGACTCTGAGACTATCGAACTCGCACCTGAATACGCTGA
GGCTATCGCAACCCGTGCAGGTGCGCTGGCTGGCATCTCTCCGATGTTCCAACCTTGCGTAGTTCCT
CCTAAGCCGTGGACTGGCATTACTGGTGGTGGCTATTGGGCTAACGGTCGTCGTCCTCTGGCGCTGG
TGCGTACTCACAGTAAGAAAGCACTGATGCGCTACGAAGACGTTTACATGCCTGAGGTGTACAAAG
CGATTAACATTGCGCAAAACACCGCATGGAAAATCAACAAGAAAGTCCTAGCGGTCGCCAACGTAA
TCACCAAGTGGAAGCATTGTCCGGTCGAGGACATCCCTGCGATTGAGCGTGAAGAACTCCCGATGA
AACCGGAAGACATCGACATGAATCCTGAGGCTCTCACCGCGTGGAGACGTGCTGCCGCTGCTGTGT
ACCGCAAGGACAAGGCTCGCAAGTCTCGCCGTATCAGCCTTGAGTTCATGCTTGAGCAAGCCAATA

AGTTTGCTAACCATAAGGCCATCTGGTTCCCTTACAACATGGACTGGCGCGGTCGTGTTTACGCTGT
GTCAATGTTCAACCCGCAAGGTAACGATATGACCAAAGGACTGCTTACGCTGGCGAAAGGTAAACC
AATCGGTAAGGAAGGTTACTACTGGCTGAAAATCCACGGTGCAAACTGTGCGGGTGTCGATAAGGT
TCCGTTCCCTGAGCGCATCAAGTTCATTGAGGAAAACCACGAGAACATCATGGCTTGCGCTAAGTCT
CCACTGGAGAACACTTGGTGGGCTGAGCAAGATTCTCCGTTCTGCTTCCTTGCGTTCTGCTTTGAGTA
CGCTGGGGTACAGCACCACGGCCTGAGCTATAACTGCTCCCTTCCGCTGGCGTTTGACGGGTCTTGC
TCTGGCATCCAGCACTTCTCCGCGATGCTCCGAGATGAGGTAGGTGGTCGCGCGGTTAACTTGCTTC
CTAGTGAAACCGTTCAGGACATCTACGGGATTGTTGCTAAGAAAGTCAACGAGATTCTACAAGCAG
ACGCAATCAATGGGACCGATAACGAAGTAGTTACCGTGACCGATGAGAACACTGGTGAAATCTCTG
AGAAAGTCAAGCTGGGCACTAAGGCACTGGCTGGTCAATGGCTGGCTTACGGTGTTACTCGCAGTG
TGACTAAGCGTTCAGTCATGACGCTGGCTTACGGGTCCAAAGAGTTCGGCTTCCGTCAACAAGTGCT
GGAAGATACCATTCAGCCAGCTATTGATTCCGGCAAGGGTCTGATGTTCACTCAGCCGAATCAGGCT
GCTGGATACATGGCTAAGCTGATTTGGGAATCTGTGAGCGTGACGGTGGTAGCTGCGGTTGAAGCA
ATGAACTGGCTTAAGTCTGCTGCTAAGCTGCTGGCTGCTGAGGTCAAAGATAAGAAGACTGGAGAG
ATTCTTCGTGAGACCGAAAGTGAAACGTGATTTCATGCGTCATTTTGAACATTTTGTAAATCTTATTT
AATAATGTGTGCGGCAATTCACATTTAATTTATGAATGTTTTCTTAACATCGCGGCAACTCAAGAAA
CGGCAGGTTCGGATCTTAGCTACTAGAGAAAGAGGAGAAATACTAGATGCGTAAAGGCGAAGAGCT
GTTCACTGGTGTCGTCCCTATTCTGGTGGAACTGGATGGTGATGTCAACGGTCATAAGTTTTCCGTGC
GTGGCGAGGGTGAAGGTGACGCAACTAATGGTAAACTGACGCTGAAGTTCATCTGTACTACTGGTA
AACTGCCGGTTCCTTGGCCGACTCTGGTAACGACGCTGACTTATGGTGTTCAGTGCTTTGCTCGTTAT
CCGGACCATATGAAGCAGCATGACTTCTTCAAGTCCGCCATGCCGGAAGGCTATGTGCAGGAACGC
ACGATTTCCTTTAAGGATGACGGCACGTACAAAACGCGTGCGGAAGTGAAATTTGAAGGCGATACC
CTGGTAAACCGCATTGAGCTGAAAGGCATTGACTTTAAAGAGGACGGCAATATCCTGGGCCATAAG
CTGGAATACAATTTTAACAGCCACAATGTTTACATCACCGCCGATAAACAAAAAAATGGCATTAAA
GCGAATTTTAAAATTCGCCACAACGTGGAGGATGGCAGCGTGCAGCTGGCTGATCACTACCAGCAA
AACACTCCAATCGGTGATGGTCCTGTTCTGCTGCCAGACAATCACTATCTGAGCACGCAAAGCGTTC
TGTCTAAAGATCCGAACGAGAAACGCGATCATATGGTTCTGCTGGAGTTCGTAACCGCAGCGGGCA
TCACGCATGGTATGGATGAACTGTACAAATGACCAGGCATCAAATAAAACGAAAGGCTCAGTCGAA
AGACTGGGCCTTTCGTTTTATCTGTTGTTTGTCGGTGAACGCTCTCTACTAGAGTCACACTGGCTCAC
CTTCGGGTGGGCCTTTCTGCGTTTATAGGTCTCACGCTGACCAGTTGCACGAGTCTCAATTGGACAA
AATGCCAGCACTTCCGGCTAAAGGTAACTTGAACCTCCGTGACATCTTAGAGTCGGACTTCGCGTTC
GCGTAATAGGATCTGCATCGCAGGATGCTGCTGGCTACCCTGTGGAACACCTACATCTGTATTAACG
AAGCGCTAACCGTTTTTATCAGGCTCTGGGAGGCAGAATAAATGATCATATCGTCAATTATTACCTC
CACGGGGAGAGCCTGAGCAAACTGGCCTCAGGCATTTGAGAAGCACACGGTCACACTGCTTCCGGT
AGTCAATAAACCGGTAAACCAGCAATAGACATAAGCGGCTATTTAACGACCCTGCCCTGAACCGAC
GACCGGGTCGAATTTGCTTTCGAATTTCTGCCATTCATCCGCTTATTATCACTTATTCAGGCGTAGCA
ACCAGGCGTTTAAGGGCACCAATAACTGCCTTAAAAAAATTACGCCCCGCCCTGCCACTCATCGCAG
TACTGTTGTAATTCATTAAGCATTCTGCCGACATGGAAGCCATCACAAACGGCATGATGAACCTGAA
TCGCCAGCGGCATCAGCACCTTGTCGCCTTGCGTATAATATTTGCCCATGGTGAAAACGGGGGCGAA
GAAGTTGTCCATATTGGCCACGTTTAAATCAAAACTGGTGAAACTCACCCAGGGATTGGCTGAGAC
GAAAAACATATTCTCAATAAACCCTTTAGGGAAATAGGCCAGGTTTTCACCGTAACACGCCACATCT
TGCGAATATATGTGTAGAAACTGCCGGAAATCGTCGTGGTATTCACTCCAGAGCGATGAAAACGTTT
CAGTTTGCTCATGGAAAACGGTGTAACAAGGGTGAACACTATCCCATATCACCAGCTCACCGTCTTT
CATTGCCATACGGAATTCCGGATGAGCATTCATCAGGCGGGCAAGAATGTGAATAAAGGCCGGATA
AAACTTGTGCTTATTTTTCTTTACGGTCTTTAAAAAGGCCGTAATATCCAGCTGAACGGTCTGGTTAT
AGGTACATTGAGCAACTGACTGAAATGCCTCAAAATGTTCTTTACGATGCCATTGGGATATATCAAC
GGTGGTATATCCAGTGATTTTTTTCTCCATTTTAGCTTCCTTAGCTCCTGAAAATCTCGATAACTCAA
AAAATACGCCCGGTAGTGATCTTATTTCATTATGGTGAAAGTTGGAACCTCTTACGTGCCGATCAAC
GTCTCATTTTCGCCAAAAGTTGGCCCAGGGCTTCCCGGTATCAACAGGGACACCAGGATTTATTTAT
TCTGCGAAGTGATCTTCCGTCACAGGTATTTATTCGGCGCAAAGTGCGTCGGGTGATGCTGCCAACT
TACTGATTTAGTGTATGATGGTGTTTTTGAGGTGCTCCAGTGGCTTCTGTTTCTATCAGCTGTCCCTC
CTGTTCAGCTACTGACGGGGTGGTGCGTAACGGCAAAAGCACCGCCGGACATCAGCGCTAGCGGAG
TGTATACTGGCTTACTATGTTGGCACTGATGAGGGTGTCAGTGAAGTGCTTCATGTGGCAGGAGAAA
AAAGGCTGCACCGGTGCGTCAGCAGAATATGTGATACAGGATATATTCCGCTTCCTCGCTCACTGAC
TCGCTACGCTCGGTCGTTCGACTGCGGCGAGCGGAAATGGCTTACGAACGGGGCGGAG
